# Xist-mediated gene silencing promotes T cell-driven autoimmunity

**DOI:** 10.64898/2026.10.15.759458

**Authors:** Nicole M. Wilkinson, Celine Park, Sumedha Chowdhury, David W. Dawson, Kathrin Plath, Maureen A. Su

**Affiliations:** Department of Microbiology, Immunology, and Molecular Genetics, David Geffen School of Medicine, University of California, Los Angeles, CA 90095; Department of Biological Chemistry, David Geffen School of Medicine, University of California, Los Angeles, CA 90095; Department of Pathology and Laboratory Medicine, Jonsson Comprehensive Cancer Center, David Geffen School of Medicine, University of California, Los Angeles, CA 90095; Department of Pediatrics, David Geffen School of Medicine, University of California, Los Angeles, CA 90095

## Abstract

Four out of five individuals affected by autoimmune disease are female. Yet, sex-specific factors that drive autoimmune predisposition in females remain poorly understood. Recent studies have implicated the female-specific long non-coding RNA Xist in autoimmunity through its ability to activate immune responses as a TLR7 ligand and through the immunogenic properties of Xist-associated proteins, stimulating dendritic cells and B cells. Here, we identify a T cell-intrinsic role for Xist in promoting autoimmunity. T cell-specific Xist deletion in mice had minimal effects on T cell populations at rest but impaired T cell fitness and increased the frequency of regulatory T cells (Tregs) in a competitive transfer setting. Xist deficiency derepressed X-linked genes, including *Foxp3*, the lineage-defining transcription factor for Tregs and *Sh2d1a*, encoding the SLAM-associated adaptor SAP, which regulates T cell activation and contraction and is linked to monogenic colitis. Finally, Xist deficiency impaired the colitogenic capacity of CD4+ T cells in an adoptive transfer model. Together, these findings reveal a previously unrecognized role for Xist-mediated gene silencing in supporting CD4+ T cell persistence and colitogenic capacity.

## INTRODUCTION

Autoimmune diseases are among the most prevalent chronic diseases in the United States^1^. Remarkably, females account for 80% of those affected by autoimmunity. Identifying sex-linked factors that modulate autoimmunity risk would therefore be a major step towards understanding pathogenesis. The female-specific long non-coding RNA (lncRNA) Xist is well-known for initiating X-chromosome inactivation (XCI) during embryogenesis^2-6^. Subsequent XCI maintenance, however, was long-considered to be independent of Xist^7^, but this has been challenged by recent reports^8-11^. In differentiated female cells, for example, higher Xist expression results in progressively greater levels of X-linked gene silencing^10^, demonstrating that Xist controls gene activity on the inactive X chromosome (Xi) beyond early development. Moreover, higher Xist expression is associated with two female-biased autoimmune conditions [systemic lupus erythematosus (SLE) and primary biliary cholangitis]^12,13^, which suggests a possible role for Xist in autoimmune predisposition. How Xist promotes autoimmunity, however, is incompletely understood.

Recent work in SLE has focused on Xist’s cell-extrinsic role in activating myeloid cells and B cells^8,11,12,14^. These studies have reported “non-canonical” roles of Xist beyond XCI, including an immunogenic function of the Xist RNA-protein complexes in inducing B cell autoantibody production^14^. Additionally, Xist RNA has been reported to function as a TLR7 agonist, leading to plasmacytoid dendritic cell activation and dysregulation^12^. In contrast to these pro-inflammatory roles, Xist in B cells has also been shown to suppress *Tlr7* expression, thereby limiting the formation of atypical B cells and autoimmune disease development. Apart from B cells and myeloid cells, CD4+ T cells also play a critical role in female-biased autoimmune disorders^15-17^. Recent reports have demonstrated that Xist localization is dynamic in CD4+ T cells upon activation and in the setting of autoimmune disease^18-20^. However, the role of Xist in CD4+ T cells, as well as its impact on T cell-mediated autoimmune conditions, is not currently known.

Here, we show that Xist in T cells promotes cellular fitness and enhances the autoimmune capacity of CD4+ T cells in a cell-intrinsic manner. Xist ablation in T cells derepressed X-linked genes such as *Foxp3* and *Sh2d1a*, whose loss-of-function has been linked to early-onset colitis^21-23^. Moreover, T cell-specific Xist-deficiency impaired the capacity of CD4+ T cells to induce colitis upon transfer into immunodeficient hosts. These findings highlight the potential of targeting Xist and/or Xist-regulated genes to limit pathogenic T cell responses in females.

## RESULTS

### Xist Localization in CD4+ T cells Is Activation State-Dependent

While recent work has demonstrated Xist’s role in the activation of B cells^8,14^ and myeloid cells^12^, its role in T cells remains unclear. This knowledge gap is particularly significant given the crucial role of CD4+ T cells in the pathogenesis of Crohn’s disease and other autoimmune conditions^24^. In support of a potential role for Xist in controlling CD4+ T cell function, Xist expression has been reported to be dynamically regulated in human and mouse T cells^18-20,25^. Consistent with this, using RNA fluorescence in situ hybridization (FISH), we detected a strong Xist signal in mouse CD4+ T cells activated with anti-CD3/anti-CD28 antibodies, whereas no signal was detectable in naïve cells **(Fig. 1A)**. XIST localization also differed between naïve and activated human CD4+ T cells, with a small punctate pattern in naïve cells, and a brighter cloud-like signal in activated CD4+ T cells **(Supp. Fig. 1A)**. Thus, Xist’s distribution changes with CD4+ T cell activation state across species, raising the possibility of a state-dependent role for Xist in T cells.

**Fig 1:**
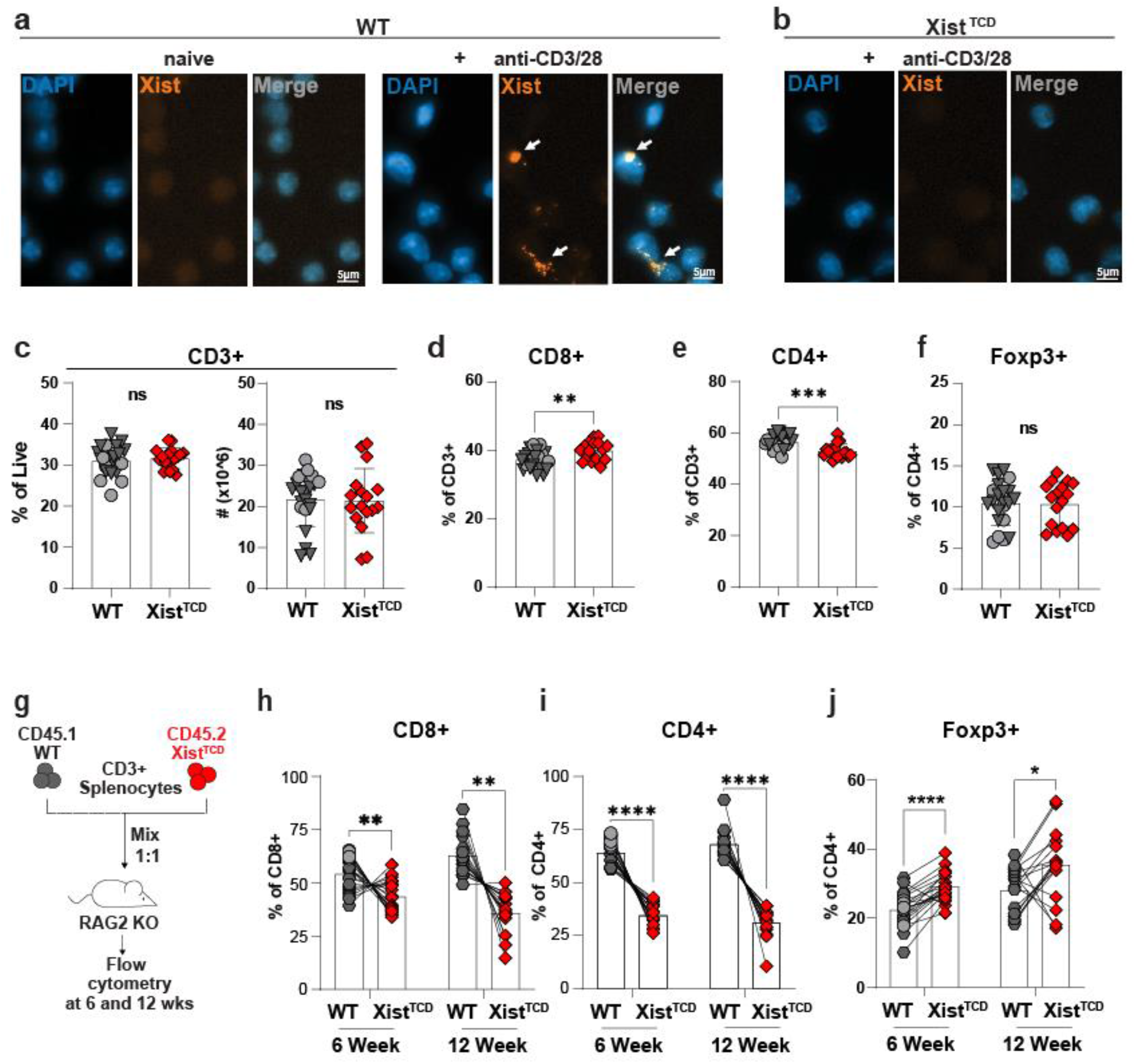
Xist minimally affects steady state T cell composition, but significantly affects expanding T cell fitness. **a-b**, Xist fluorescent in situ hybridization (FISH) in naïve WT**(a, left)**, activated (48 hours, 1ug/mL anti-CD3/28) WT **(a, right)**, and activated Xist^TCD^ **(b)** CD4+ T cells. **c-f**, Flow cytometry of splenocytes from 8-week WT (n = 22) and Xist^TCD^ (n = 17) animals. Percentage **(c, left)** and absolute number **(c, right)** CD3+ splenocytes of live splenocytes. Percentage CD8+ **(d)** and CD4+ **(e)** T cells of CD3+ splenocytes. Percentage Foxp3+ T cells **(f)** among CD3+CD4+ splenocytes. **g,** Schematic of CD3+ T cell co-transfer [1:1 WT (CD45.1) and Xist^TCD^ (CD45.2)] into RAG2 KO recipients for flow cytometry. **h-i,** Relative frequency WT and Xist^TCD^ donor cells within CD8+ **(h)** and CD4+ **(i)** T cells 6 (n = 20) and 12 (n = 17) weeks post-transfer. **j**, Percentage Foxp3+ T cells of WT and Xist^TCD^ CD3+CD4+ splenocytes 6 and 12 weeks post-transfer. Data are representative of 2-5 experiments. Samples were compared using unpaired two-tailed Student’s *t*-test **(c-f)** or paired two-tailed Student’s *t*-test **(h-j).** Error bars represent standard deviation. \**P* < 0.05; \*\**P* < 0.01; \*\*\**P* < 0.001; \*\*\*\**P* < 0.0001. For **(c-f)** and **(h-j)**, =CD4-Cre negative Xist^fl/fl^, =CD4-Cre negative Xist^wt/wt^, =CD4-Cre positive Xist^wt/wt^ mice.

### Xist Minimally Perturbs Peripheral T cell Subset Distribution under Steady-State Conditions

To define the role of Xist in T cells, we generated C57BL/6 mice with T cell-specific Xist deficiency (CD4-cre+ Xist^fl/fl^, herein referred to as Xist^T^ ^Cell^ ^Deficient^ or Xist^TCD^ mice). Xist RNA FISH confirmed that Xist is present in activated T cells from wildtype mice [Cre-negative (CD4-cre-Xist^fl/fl^), hereafter termed WT mice], but absent in T cells from Xist^TCD^ mice **(Fig. 1A, right, and 1B)**. The CD4-cre transgene drives Xist deletion when thymocytes first upregulate CD4 at the double positive (DP) stage of development^26^. Thus, we evaluated thymic composition in 8-week-old WT and Xist^TCD^ mice. No differences were seen in the frequency or absolute numbers of double positive (DP; CD4+ CD8+), CD4+ single positive (CD4+ SP), or CD8+ single positive (CD8+ SP) thymocytes **(Supp. Fig. 2A-C),** suggesting that Xist loss does not alter thymocyte subset distribution.

We next evaluated the effect of T cell-specific Xist deficiency on peripheral T cell numbers. Under steady state conditions, splenic CD3+ T cell frequency and absolute numbers were grossly unchanged between 8-week-old WT and Xist^TCD^ mice **(Fig. 1C; Supp. Fig. 2D)**. However, the composition of the CD3+ population in Xist^TCD^ mice was modestly but significantly altered, with increased frequency of CD8+ and decreased frequency of CD4+ T cells **(Fig. 1D,E; Supp. Fig. 2E).** No differences, however, were observed in the absolute numbers of CD3+ CD8+ or CD3+ CD4+ T cells **(Supp. Fig. 2F).** Within CD4+ T cells, no differences were seen in the frequency of FOXP3+ (Treg) **(Fig. 1F; Supp. Fig. 2G),** IFNy-producing (Th1) or IL17A-producing (Th17) CD4+ T helper subsets **(Supp. Fig. 2H,I)**. Thus, in the periphery of unperturbed mice, Xist deficiency is associated with slight changes in CD8+ and CD4+ T cell frequencies but does not alter the total number of CD3+ T cells, absolute numbers of CD8+ and CD4+ T cells, or the frequencies of functional CD4+ T cell subsets.

### Xist Maintains Repression of X-linked ‘Escapees’ in Steady-State CD4+ T Cells

Given Xist’s role in inactivating X-linked genes in B cells and other cell types^5,8,27^, we sought to determine how T cell-specific Xist deficiency alters the transcriptome. Although the frequencies of CD8+ and CD4+ T cell were both altered in Xist^TCD^ spleens (**Fig. 1D,E**), we focused on CD4+ T cells because of their importance in female-biased autoimmunity^15-17^. Bulk RNA-sequencing (RNA-seq) of splenic CD4+ T cells from 8-week-old WT and Xist^TCD^ mice (**Supp. Fig. 3A**) revealed 396 differentially expressed genes, of which 20 were X-linked (**Supp. Fig. 3B**). Of these 20 genes, the majority (15) were upregulated in Xist^TCD^ CD4+ T cells **(Supp. Fig. 3B,C; Supp. Table 1)**, consistent with the loss of Xist-dependent silencing. We tested whether ‘Xist-dependent’ genes (defined as X-linked genes upregulated upon Xist deletion) exhibit epigenetic or genomic features that may explain their continued reliance on Xist. H3K27me3 levels^28^, DNA methylation^29^, and X chromosome position (distance from the Xist locus) did not significantly differ between Xist-dependent and Xist-independent X-linked genes **(Supp. Fig 3D-F)**. Xist-dependent genes were, however, enriched for immune genes such as *Cxcr3, Firre,* and *Foxp3*, compared to Xist-independent genes (OR = 5.696, *P* =0.025, Fisher’s exact test) **(Supp. Fig. 3G, Supp. Table 2)**.

In B cells and other cell types, Xist loss has been reported to preferentially derepress genes that escape XCI (‘escapees’)^8,30^. These ‘genes are not fully silenced and exhibit expression from the Xi in wildtype cells. To test whether ‘escapees’ are preferentially derepressed following Xist loss in CD4+ T cells, we compared the frequency of escapees among the 15 Xist-dependent genes and Xist-independent X-linked genes. Escape genes, as defined by Hauth et al^10^, were over-represented among Xist-dependent genes **(Supp. Fig. 3H, Supp. Table 3)**, comprising 47% of Xist-dependent genes compared to 19% of Xist-independent genes (OR = 3.811, *P* = 0.013, Fisher’s exact test). Thus, under steady-state conditions, Xist preferentially maintains repression of escape genes in CD4+ T cells.

### During Homeostatic Expansion, T cell-specific Xist Promotes CD4+ T Cell Fitness and Limits Treg Frequencies

T cell homeostatic expansion, in which T cells expand to restore T cell numbers in the setting of lymphopenia, is a cytokine- and self-antigen-driven process^31,32^. Importantly, homeostatic expansion has been linked to autoimmunity predisposition^33^, as highlighted by the frequent association between primary immunodeficiency syndromes and the development of autoimmune diseases^34^ such as inflammatory bowel disease^35,36^. To examine Xist’s function in T cells undergoing homeostatic expansion, we evaluated Xist-deficient T cells after transfer into lymphopenic hosts. Congenically marked WT and Xist^TCD^ CD3+ T cells were mixed at a 1:1 ratio for adoptive transfer into immunodeficient RAG2 KO hosts **(Fig. 1G)**. Six weeks after transfer, the proportion of Xist^TCD^ donor cells was significantly reduced among both CD4+ and CD8+ T cell populations in recipient mice **(Fig. 1H,I)**, with Xist-deficient T cells comprising ∼35% of all CD4+ T cells and ∼43% of all CD8+ T cells. Twelve weeks after transfer, Xist^TCD^ donor cells comprised ∼31% of CD4+ T cells and ∼38% of CD8+ T cells in host spleens **(Fig. 1H,I)**. Thus, in the setting of lymphopenia, T cell Xist deficiency is associated with reduced competitive fitness, with a more pronounced disadvantage among CD4+ than CD8+ T cells. Among CD4+ T cells, Foxp3+ Tregs derived from Xist^TCD^ donors were significantly increased in frequency at both 6 and 12 weeks after transfer **(Fig. 1J).** This finding is consistent with the observation that *Foxp3* transcript levels are upregulated in Xist^TCD^ in steady-state conditions **(Supp. Fig. 3C)**.

Of note, similar results were seen when comparisons were made with Cre-positive mice carrying wildtype Xist alleles (CD4-cre+ Xist^wt/wt^) as controls, indicating that our findings are not merely due to Cre expression and potential Cre toxicity **(Fig. 1C-F; 1H-J)**^37^. Taken together, our findings suggest a key role for Xist in supporting CD4+ T cell competitive fitness while limiting the frequency of Foxp3+ Tregs during homeostatic expansion.

### In CD4+ T cells Undergoing Homeostatic Expansion, Xist Silences *Sh2d1a/*SAP*, Foxp3*, and other X-linked Immune Genes

To identify Xist-regulated transcriptomic changes during homeostatic expansion, we performed RNA-seq on WT and Xist^TCD^ CD4+ T cells at six and twelve weeks after CD4+ T cell transfer (**Fig. 2A,B**). Compared to T cells under steady-state conditions, an increased number of X-linked genes were upregulated in Xist-deficient T cells undergoing homeostatic expansion. While only 15 X-linked genes were upregulated at steady state (**Supp. Fig. 3B,C**), 49 were upregulated at 6 weeks and 68 were upregulated at 12 weeks after transfer **(Fig. 2C,D; Supp. Table 1)**. Comparison of differentially expressed genes (DEGs) in CD4+ T cells at 6 and 12 weeks after transfer revealed a high degree of overlap, with 41 shared upregulated X-linked genes (*P* = 1.63e-49, Fisher’s exact test). These 41 shared genes were used to define an Xist-dependent signature in activated CD4+ T cells (hereafter the ‘T_Act_ signature’) **(Fig. 2E)**. While no differences were seen in DNA methylation^29^ or X chromosome position (distance from the Xist locus) **(Supp. Fig. 4B,C),** ‘T_Act_ signature’ genes showed higher levels of H3K27me3^28^ compared to remaining X-linked genes **(Supp. Fig. 4A)**. This is in contrast to Xist-dependent genes in B cells^8^, which have been reported to have lower H3K27me3 levels at Xist-dependent gene bodies. Xist deficiency derepresses a small subset of genes in B cells^38^ and hematopoietic stem cells [LSK+ (Lin-, Sca-1+, c-Kit+) and LSK- (Lin-, Sca-1, c-Kit)]^30^ that are shared with the ‘T_Act_ signature’ **(Supp. Fig. 4D-F, Supp. Table 4)**. However, most ‘T_Act_ signature’ genes are distinct from those upregulated by Xist deficiency in other immune cell types. Together, these analyses suggest that the repertoire of Xist-dependent genes differs across immune cell types.

**Fig 2:**
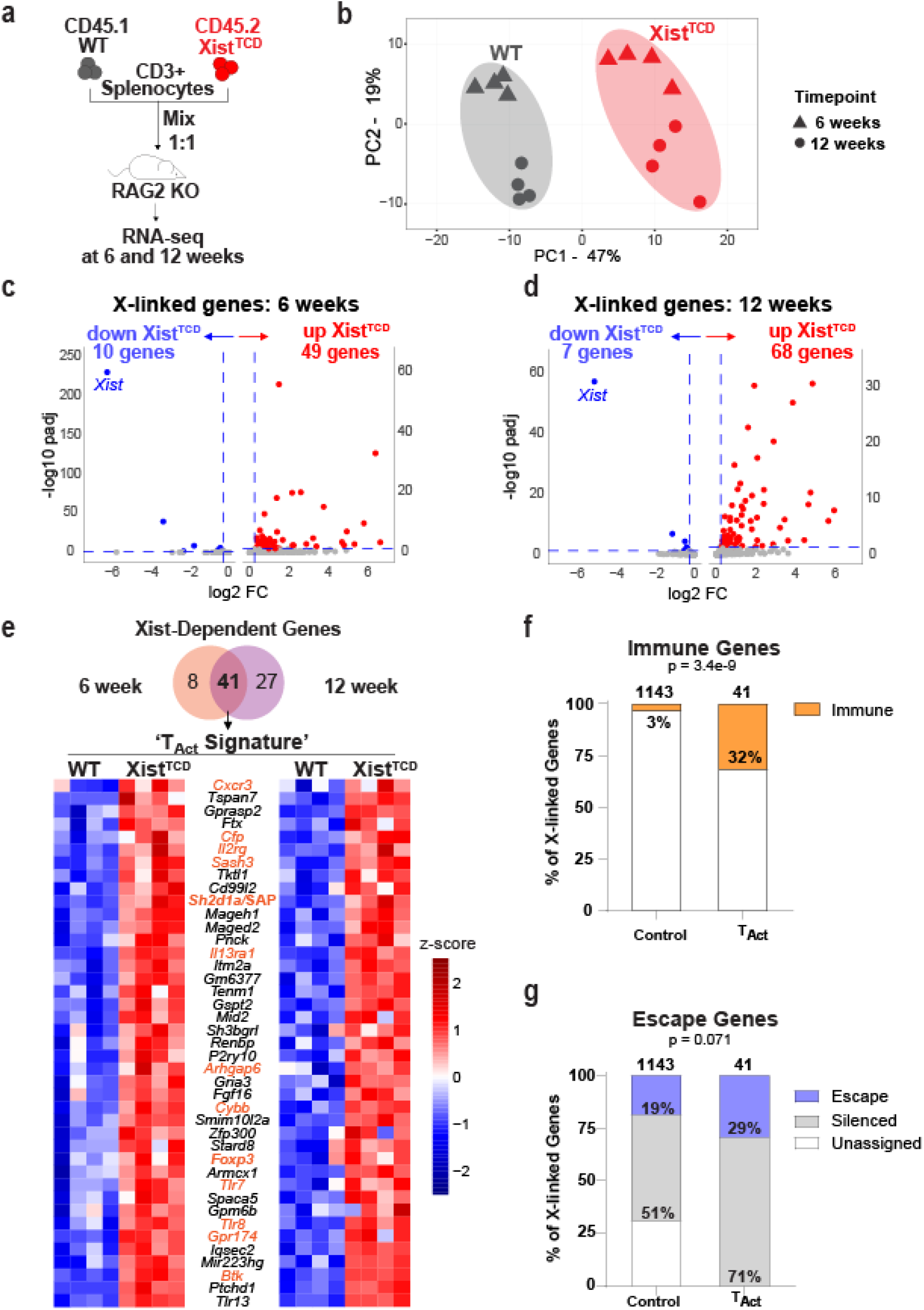
In CD4+ T cells undergoing homeostatic expansion, T cell Xist silences *Sh2d1a*/SAP, *Foxp3* and other X-linked immune genes, **a**, Experimental set-up showing CD3+ T cell co-transfer [1:1 WT (CD45.1) and Xist^TCD^ (CD45.2)] into RAG2 KO recipients for RNAseq of flow-sorted WT or Xist^TCD^ CD4+ T cells from host spleens. **b,** Principal component analysis of WT and Xist^TCD^ CD4+ T cells 6 weeks and 12 weeks after adoptive transfer (n = 4 per group, samples represent biologic replicates). **c-d**, Volcano plot of X-linked genes at the 6-week **(c)** and 12-week **(d)** timepoint. **e,** Heatmap of Xist-dependent genes (X-linked, padj < 0.05, log2FC > 0.25) common to the 6- and 12-week time points. Immune-related genes shown in orange. **f-g,** Percentage immune genes **(f)** and escape or silenced genes **(g)** among T_Act_ Signature genes compared to control (remaining X-linked genes). Enrichment of immune-related genes **(f)** and escape genes **(g)** was assessed using Fisher’s exact test.

Immune-related genes were overrepresented among ‘T_Act_ signature’ genes, making up 32% of ‘T_Act_ signature’ genes compared to 3% of other X-linked genes (OR = 13.067, *P* = 3.4e-9, Fisher’s exact test) **(Fig. 2F, Supp. Table 2)**. These genes included immunoregulatory factors such as *Sh2d1a/*SAP and *Foxp3*. Among the ‘T_Act_ signature’ genes, 29% were escapees, representing a non-significant enrichment compared to other X-linked genes [19%; OR = 1.806, *P* = 0.071, Fisher’s exact test] **(Fig. 2G, Supp. Table 3)**. This represents a lower percentage than that observed in steady-state CD4+ T cells (47%) **(Supp. Fig. 3H).** Thus, Xist-dependent repression during homeostatic expansion extends beyond known escape genes to include genes normally subject to XCI.

### Xist-Deficient T Cells Have An Increased Susceptibility to Apoptosis and Overexpress *Sh2d1a*/SAP

We next sought to define cellular mechanisms underlying the fitness defect in Xist-deficient CD4+ T cells. One possibility is that Xist deficiency impairs CD4+ T cell proliferation. However, no differences were observed in the expression of the proliferation marker Ki67 at 6 or 12 weeks after transfer into lymphopenic recipients **(Fig. 3A)**. Further, no difference was seen in Ki67 expression five hours after *in vivo* anti-CD3 stimulation **(Fig. 3B)**. Another possibility is that Xist deficiency increases apoptosis. Since apoptotic cells are subject to rapid phagocytosis by macrophages^39^, we evaluated markers of. Apoptosis shortly following *in vivo* activation. WT and Xist^TCD^ CD4+ T cells were assessed by flow cytometry five hours after T cell receptor (TCR) stimulation with anti-CD3 antibody injection^40^. An increased frequency of Annexin V-positive cells was observed in Xist^TCD^ CD4+ T cells compared to WT at this timepoint **(Fig. 3C)**. Consistent with these findings, Xist^TCD^ CD4+ T cells showed a trend toward an increased frequency of Annexin V-positive cells and a significantly increased frequency of cleaved caspase 3-positive cells after in vitro anti-CD3 and anti-CD28 antibody stimulation (**Fig. 3D,E**). Similarly, an increased frequency of Annexin V-positive cells was noted in human CD4+ T cells after CRISPR-Cas9-mediated XIST deletion **(Fig. 3F; Supp. Fig. 5A,B)**. Together, these findings suggest that the reduced fitness of Xist^TCD^ CD4+ T cells is associated with increased propensity to apoptosis.

**Fig 3:**
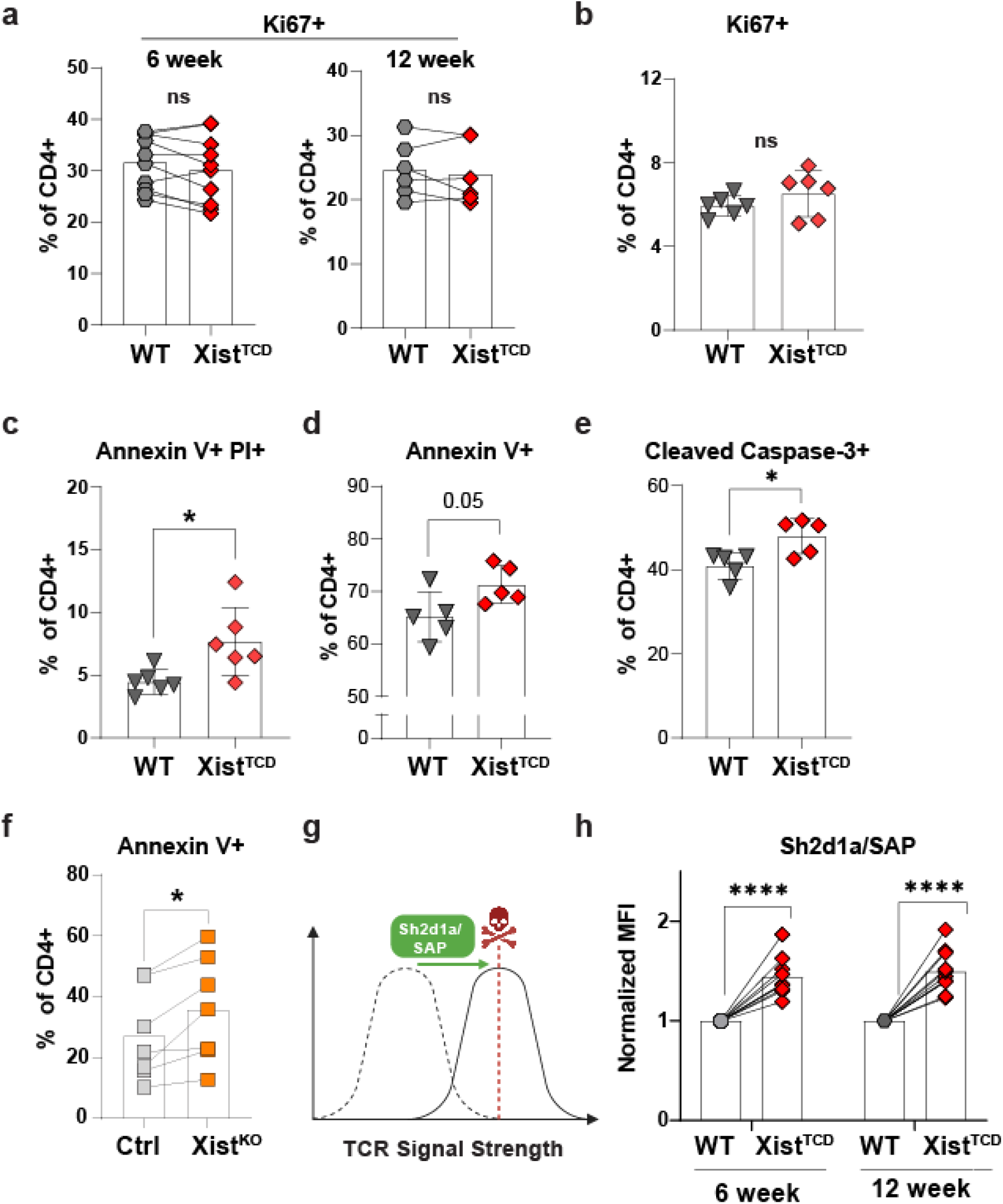
Xist-deficient CD4+ T cells are more prone to apoptosis and overexpress *Sh2d1a*/SAP. **a**, Percentage Ki67+ of WT and Xist^TCD^ CD4+ T cells in RAG2 KO recipient spleens 6 (n = 10) and 12 (n = 6) weeks following CD3+ T cell co-transfer (1:1). **b-c,** Percentage Ki67+ **(b)** and Annexin V+ PI+ **(c)** of CD4+ splenocytes from WT (n = 6) and Xist^TCD^ (n = 6) mice treated with anti-CD3 antibodies (10 ug i.v., 5 hours). **d-e**, Percentage annexin V+ **(d)** and cleaved caspase-3+ **(e)** of WT (n = 5) and Xist^TCD^ (n = 5) CD4+ splenocytes following in vitro T cell stimulation with plate-bound anti-CD3/28 antibodies (1ug/mL) **f,** Percentage annexin V+ of control or Xist^KO^ (CRISPR-edited) human primary CD4+ T cells (Donor 1: n = 3, Donor 2: n = 2, Donor 3: n = 2) following initial culture with plate-bound anti-CD3/28 (1ug/mL), rest (hIL2, 50U/mL), and restimulation with soluble anti-CD3 (1ug/mL). **g,** Schematic of Sh2d1a/SAP role in activation induced cell death. Adapted from Snow et al (2009). **(h)** SAP MFI (normalized to WT) in WT and Xist^TCD^ CD4+ T cells from RAG2KO recipient spleens 6 (n = 10) and 12 (n = 11) weeks following CD3+ T cell co-transfer (1:1). Data are representative of 2 – 4 experiments. Samples were compared using unpaired two-tailed Student’s *t*-test **(b-e)** or paired two-tailed Student’s *t*-test **(a,f,h).** Error bars represent standard deviation. \**P* < 0.05; \*\*\*\**P* < 0.0001.

Based on these findings, we hypothesized that Xist deficiency may derepress X-linked genes that promote T cell apoptosis. Among X-linked genes in the ‘T_Act_ signature’ is *Sh2d1a*, which encodes SAP **(Fig. 2E)**, an intracellular adaptor for the SLAM family of receptors that plays a key role in T cell apoptosis^41^ (**Fig. 3G**). Through potentiation of TCR signal strength, SAP drives T cells towards an activation threshold at which apoptosis is induced^32,33^. In patients, *SH2D1A*/SAP deficiency results in X-linked Lymphoproliferative Disorder (XLD)^26–29^, a condition associated with a failure of T cell contraction after immune challenge^42^. Thus, we hypothesized that increased SAP expression may contribute to the increased propensity for apoptosis. Indeed, flow cytometric analysis revealed elevated *Sh2d1a*/SAP protein expression in Xist^TCD^ CD4+ T cells compared to WT at 6 and 12 weeks after transfer into lymphopenic hosts (**Fig. 3H**). Thus, Xist^TCD^ T cells exhibit higher *Sh2d1a*/SAP expression during homeostatic expansion, consistent with a potential role for *Sh2d1a*/SAP in their increased susceptibility to apoptosis.

### *Sh2d1a*/SAP Overexpression Recapitulates Enhanced TCR Signaling and Apoptosis Observed Following Xist Deficiency in CD4+ T Cells

Given our data linking Xist deficiency to increased SAP expression in CD4+ T cells **(Fig. 3H)**, we next sought to determine whether increased SAP expression in CD4+ T cells is sufficient to drive a fitness deficit. Because SAP modulates signaling cascades downstream of T cell receptor (TCR) engagement^30,31^, we utilized the Nur77-GFP reporter system^43^ to define the effects of SAP overexpression on TCR signal strength. Nur77 is an immediate early gene induced proportionally to TCR signal strength, so that greater Nur77 reporter expression signifies higher TCR signal strength^43^. CD4+ T cells from Nur77-GFP mice were transduced with a SAP-overexpression retrovirus (MSCV-*Sh2d1a*-IRES-mCherry) or an empty retrovirus (MSCV-IRES-mCherry). SAP-overexpression was confirmed by flow cytometry **(Supp. Fig. 5C,D)**. We observed greater Nur77-GFP expression in SAP-overexpressing compared to control-transduced CD4+ T cells at 1 and 6 hours after TCR restimulation **(Fig. 4A,B)**. In addition to Nur77 expression, Zap70 phosphorylation at Tyr319 is induced following TCR engagement and is critical for downstream signaling cascades. An increased pZap70:Zap70 ratio is therefore indicative of greater TCR signaling^44^. We observed an increased pZap70:Zap70 ratio in SAP-overexpressing versus control-transduced CD4+ T cells **(Fig 4C; Supp. Fig. 6A**), further supporting increased TCR signaling with SAP overexpression.

**Fig 4:**
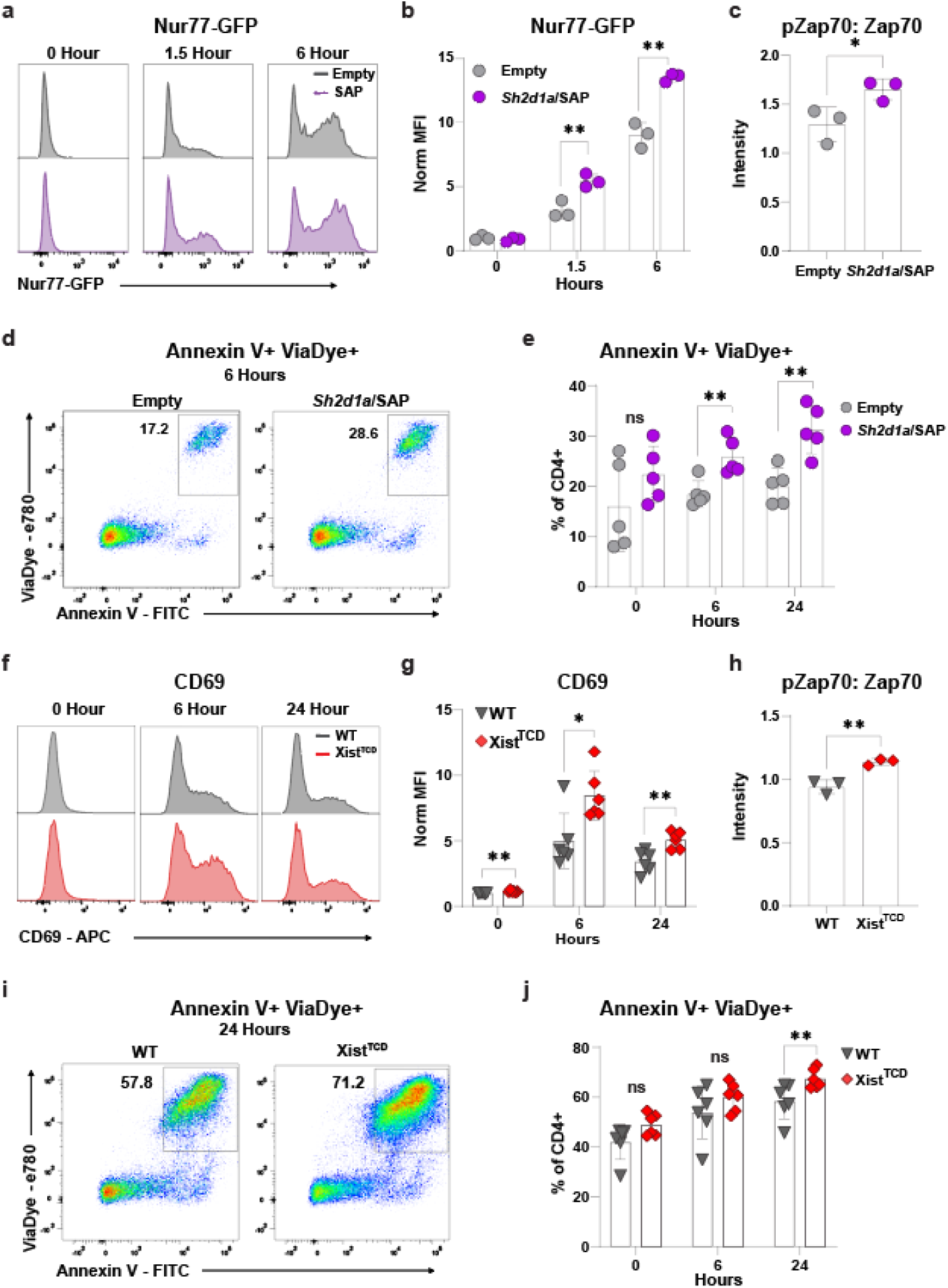
S*h*2d1a/SAP overexpression phenocopies enhanced activation and apoptosis seen with Xist deficiency in CD4+ T Cells. For all experiments, CD4+ T cells were activated with plate bound anti-CD3/28 antibodies (1ug/mL), rested in hIL2 (50IU/mL), and restimulated for assessment of activation and apoptosis. **a-b,** Representative Nur77-GFP histograms **(a)** and Nur77-GFP MFI (normalized to WT) **(b)** in SAP-overexpressing (MSCV-Shd1a-IRES-mCherry) (n = 3) and control-transduced (MSCV-Empty-IRES-mCherry) (n = 3) Nur77-GFP CD4+ T cells after soluble anti-CD3 antibody (1ug/mL) restimulation. **c,** pZap70:Zap70 western blot quantification in SAP-overexpressing (n = 3) and control-transduced (n = 3) WT CD4+ T cells after plate-bound anti-CD3/28 antibody (1ug/mL) restimulation. **d-e,** Representative dot plots **(d)** and percentage Annexin V+ ViaDye+ **(e)** of SAP-overexpressing (n = 5) and control-transduced (n = 5) WT CD4+ T cells after soluble anti-CD3 antibody (1ug/mL) restimulation. **f**-**g,** Representative CD69 histograms **(f)** and MFI (normalized to WT) **(g)** in WT (n = 6) and Xist^TCD^ (n = 6) CD4+ T cells after soluble anti-CD3 (1ug/mL) restimulation**. h,** pZAp70:Zap70 western blot quantification after plate-bound anti-CD3/28 antibody (1ug/mL) restimulation. **i-j**, Representative dot plot **(i)** and percentage Annexin V+ ViaDye+ **(j)** of WT (n = 6) and Xist^TCD^ (n = 6) CD4+ T cells after soluble anti-CD3 antibody (1ug/mL) restimulation. Samples were compared using unpaired two-tailed Student’s *t*-test (**b-c; e; g-h; j**). Error bars represent standard deviation. \**P* < 0.05; \*\**P* < 0.01; \*\*\**P* < 0.001.

Through potentiation of TCR signal strength, SAP drives T cells towards an activation threshold at which apoptosis is induced^32,33^ **(Fig. 3G)**. Thus, we next sought to determine the effects of SAP overexpression on apoptosis. To do this, we compared Annexin V expression at 6 and 24 hours after TCR restimulation. At both time points, significantly higher Annexin V levels were observed in SAP-overexpressing compared to control-transduced CD4+ T cells **(Fig. 4D,E)**. These findings support prior work^42^ demonstrating a pro-apoptotic role for SAP in T cells. We next asked, whether Xist-deficient CD4+ T cells similarly exhibit enhanced TCR signaling and apoptosis. Increased TCR activation, as evidenced by higher CD69 expression **(Fig. 4F,G)** and increased pZap70:Zap70 ratios **(Fig. 4H; Supp. Fig. 6B),** was observed following Xist depletion. Additionally, increased Annexin V staining **(Fig. 4I,J)**, indicative of higher apoptotic propensity, was detected in Xist^TCD^ CD4+ T cells. Thus, Xist^TCD^ CD4+ T cells phenocopy SAP-overexpressing CD4+ T cells, suggesting that increased SAP expression may contribute to the enhanced TCR signaling and apoptosis observed following Xist deletion.

### T cell Xist Promotes CD4+ T Cell Fitness in Autoimmune Colitis

Crohn’s disease is a female-biased and T cell-mediated autoimmune condition characterized by inflammation of the gastrointestinal tract^45-47^. Intriguingly, *SH2D1A*/SAP polymorphisms have been linked to inflammatory bowel disease^48,49^ and loss-of-function *SH2D1A*/SAP mutations result in a hyperinflammatory state associated with early-onset colitis^21,22^. At the same time, loss-of-function *FOXP3* mutations characteristically result in autoimmune colitis in infancy^23^. Thus, we hypothesized that increased SAP or *Foxp3* expression in Xist^TCD^ CD4+ T cells may suppress their colitogenic capacity.

To test this hypothesis, we utilized the CD45RB^hi^ CD4+ T cell adoptive transfer mouse model of inflammatory bowel disease^50^, in which CD45RB^hi^ CD4+ T cells are transferred into RAG2 KO recipients to induce colitis **(Supp. Fig. 5E).** We first sought to compare the fitness of colitogenic CD45RB^hi^ CD4+ T cells from WT and Xist^TCD^ mice. To control internally for levels of inflammation and other microenvironmental differences, RAG2 KO hosts were transferred with a 1:1 WT : Xist^TCD^ mix of CD45RBhi CD4+ T cells **(Fig. 5A, Supp. Fig. 5F)**. At 5-7 weeks after transfer, the proportion of Xist-deficient T cells was ∼7-fold lower than that of WT cells across secondary lymphoid tissues [spleen, mesenteric lymph node (mLN)] and colon **(Fig. 5B).** Moreover, a significant increase in the frequency of Foxp3+ Tregs was seen among Xist^TCD^ compared to WT CD4+ T cells in mLN (**Fig. 5C**). No differences were seen in the frequency of Foxp3+ Tregs in the spleen or colon **(Supp. Fig. 7A),** and no differences were seen in the frequencies of IL17-producing, IFN-g producing or IFN-g/IL17 co-producing CD4+ T cells across the tissues examined (**Supp. Fig. 7B-D**). These findings demonstrate that T cell Xist promotes the competitive fitness of colitogenic CD4+ T cells and limits the relative abundance of Foxp3+ Tregs in the mLN.

**Fig 5:**
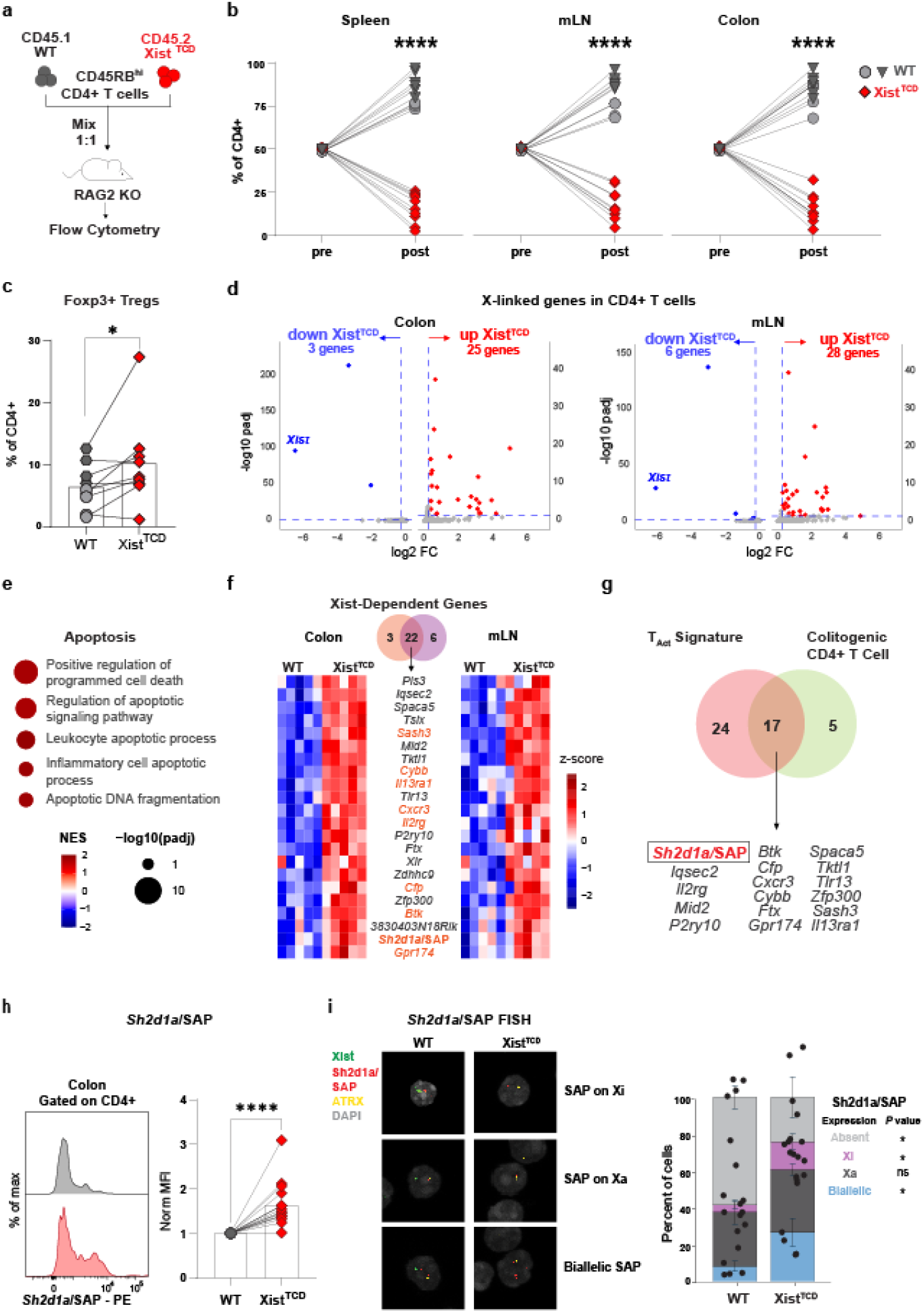
T cell Xist promotes CD4+ T cell fitness in autoimmune colitis and silences *Sh2d1a*/SAP expression on the Xi. **a**, Experiment schematic of CD45RB^hi^ CD4+ T cell co-transfer [1:1 WT (CD45.1) and Xist^TCD^ (CD45.2)] into RAG2 KO recipients for flow cytometry. **b**, Relative frequency WT and Xist^TCD^ of total CD4+ T cells in the spleen, mLN, and colon 7-9 weeks post transfer (n = 10). **c**, Percentage Foxp3+ **(c)** of WT and Xist^TCD^ CD3+CD4+ T cells in the mLN of RAG2 KO recipients (n = 9). **d-g,** WT and Xist^TCD^ CD4+ T cells were flow sorted from RAG2 KO recipients (n = 5) colon and mLN for RNA-seq 7 weeks after CD45RB^hi^ CD4+ T cell co-transfer [1:9 WT (CD45.1) and Xist^TCD^ (CD45.2), to ensure adequate numbers of Xist-deficient cells]. **d,** Volcano plots of X-linked genes in colon and mLN. **e,** Gene set enrichment analysis (GSEA) in colitogenic CD4+ T cells. **f,** Heatmap of Xist-dependent genes common to colon and mLN. Immune-related genes shown in orange. **g**, Comparison of T_Act_ Signature genes and Xist-dependent genes in colitogenic CD4+ T cells **h**, Representative Sh2d1a/SAP histograms and normalized SAP MFI of WT and Xist^TCD^ CD4+ T cells from the colon of RAG2 KO mice (n = 17) with colitis. **i,** Representative images (left) and quantification (right) of nascent transcript FISH for *Sh2d1a* in WT (n = 5) and Xist^TCD^ (n = 5) CD4+ T cells from the mLN or RAG2 KO mice with colitis. Samples were compared using paired two-tailed Student’s *t-*test (**b-c;e;h**) and Wilcoxon rank-sum test (**i**). \**P* < 0.05; \*\*\*\**P* < 0.0001.

### In Colitogenic CD4+ T Cells, Xist Maintains Repression of *Sh2d1a* on the Xi

We next sought to delineate Xist-dependent transcriptional changes in colitogenic T cells, utilizing co-transfer of WT and Xist^TCD^ CD45RB^hi^ CD4+ T cells into RAG2 KO recipients. Seven weeks after transfer, CD4+ T cells from the colon and mLN were sorted for RNA-sequencing. Of the X-linked DEGs, 25 genes were upregulated in the colon (**Fig. 5D, left**) and 28 genes were upregulated in the mLN (**Fig. 5D, right**). Despite our data suggesting that Xist preferentially represses XCI ‘escapees’ in steady-state CD4+ T cells (**Supp. Fig. 3H**), escape genes were not significantly enriched among Xist-dependent genes in colitogenic CD4+ T cells **(Supp. Fig. 8A, Supp. Table 3).** Thus, whether XCI ‘escapees’ are preferentially targeted by Xist is T cell state-dependent.

Pathway analysis revealed enrichment of gene sets relevant to T cell activation and apoptosis **(Fig. 5E, Supp. Fig. 8B)**, including “Alpha Beta T cell activation”, “Antigen receptor mediated signaling pathway”, “Positive regulation of programmed cell death”, “Regulation of apoptotic signaling pathway”, and “Leukocyte apoptotic process”. This result is consistent with enhanced TCR signaling and increased apoptotic predisposition in Xist-deficient CD4+ T cells.

Comparison of X-linked genes upregulated in Xist^TCD^ CD4+ T cells from the colon and mLN identified 22 shared genes, including *Sh2d1a* and other immune genes **(Fig. 5F)**. Despite our data showing Foxp3+ Tregs were increased in frequency in the mLN (**Fig. 5C**), Foxp3 transcript levels were not upregulated in Xist^TCD^ CD4+ T cells from the colon or mLN **(Supp. Table 1).** A caveat with this analysis, however, is that the low frequency of Foxp3+ T cells among CD4+ T cells in this model may preclude detection of subtle changes in Foxp3 expression. *Sh2d1a,* on the other hand, was one of the 22 shared, upregulated X-linked genes **(Fig. 5F)**.Of note, *Sh2d1a* was among the 17 shared X-linked genes (between colon and mLN CD4+ T cells) that were also part of the ‘T_Act_’ core signature (**Fig. 5G**). Thus, *Sh2d1a* upregulation is observed in Xist-deficient CD4+ T cells during both homeostatic expansion and colitis induction. Flow cytometric analysis verified upregulation of SAP protein in CD4+ T cells from the inflamed colon as well as the mLN (**Fig. 5H, Supp. Fig. 8C**).

To define the allelic specificity of *Sh2d1a/*SAP upregulation, we performed nascent transcript RNA FISH in CD4+ T cells sorted from the mLN of RAG2 KO recipients reconstituted with CD45RBhi CD4+ T cells. The overall frequency of *Sh2d1a*-expressing cells was increased in Xist^TCD^ CD4+ T cells compared to WT **(Fig. 5I)**. Notably, the frequency of cells exhibiting *Sh2d1a* transcription from the Xi (biallelic or Xi-only expression) increased following Xist deletion with 42% of Xist^TCD^ CD4+ T cells showing Xi expression compared to only 12% of WT cells **(Fig. 5I)**. Together, these findings demonstrate that Xist is required for the ongoing repression of *Sh2d1a* on the Xi.

### Xist Promotes CD4+ T Cells Colitogenic Capacity

Our data suggest that Xist promotes the colitogenic capacity of CD4+ T cells through the silencing of immunomodulatory genes. In support of this possibility, significantly higher XIST levels were observed in CD4+ T cells from female Crohn’s disease patients compared to healthy controls^51^ **(Fig. 6A, left)**. Importantly, no differences were seen in XIST levels between non-inflamed and inflamed sites **(Fig. 6A, right)**, suggesting that increased XIST expression is not merely a secondary consequence of local inflammation.

**Fig 6:**
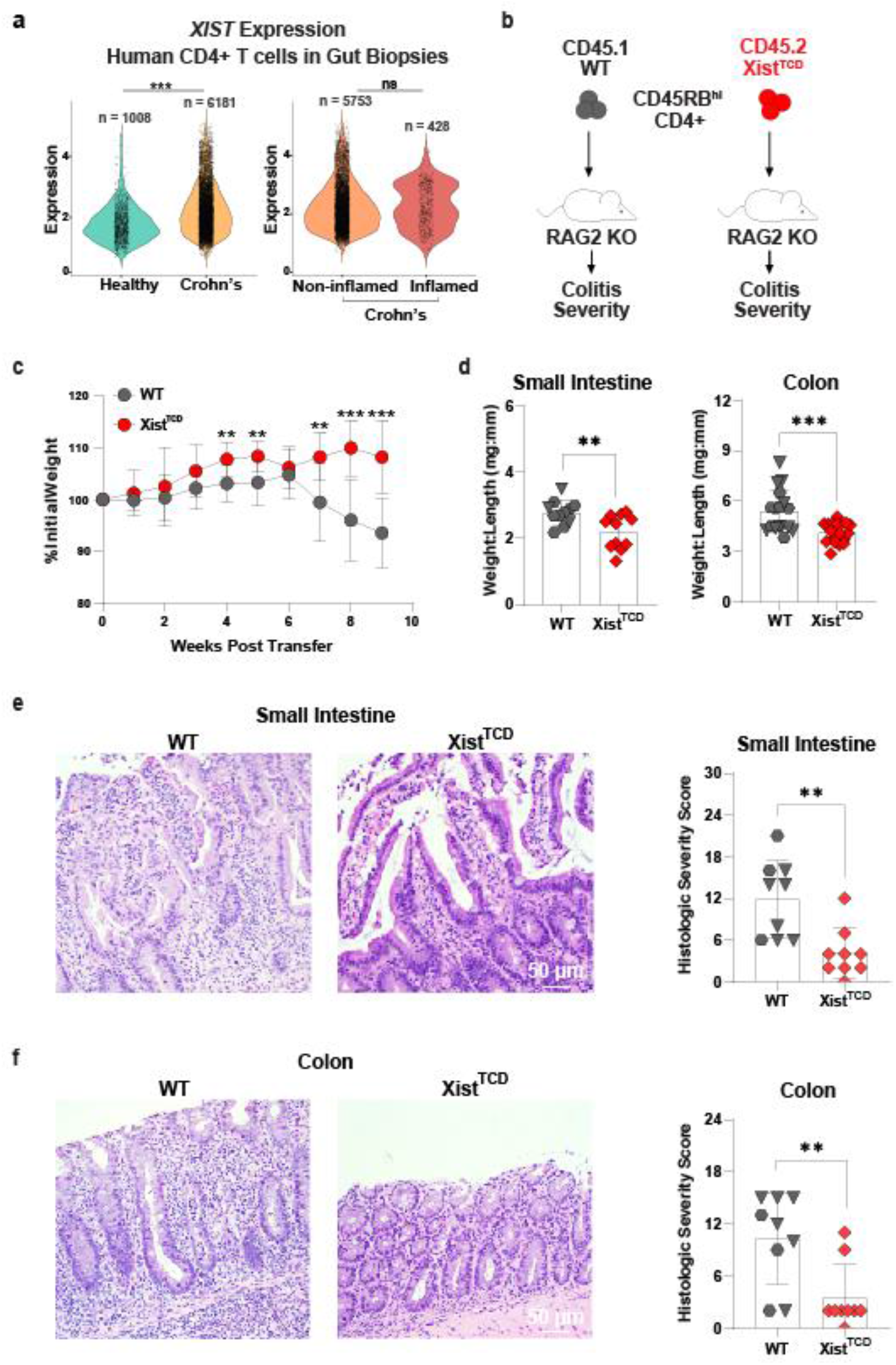
Xist promotes CD4+ T cell colitogenic capacity. **a**, Single cell RNA sequencing of CD4+ T cells in gut biopsies from females with Crohn’s Disease (CD) (n = 11) and healthy controls (n = 6). Xist expression in healthy controls vs CD (independent of inflammatory status) **(a, left)** and non-inflamed vs inflamed CD samples **(a, right)**. **b,** Colitis was induced by adoptive transfer of WT or Xist^TCD^ CD45RB hi CD4+ T cells into RAG2 KO recipients**. c,** Body weight curves for WT (n = 11) and Xist^TCD^ (n = 11) animals with colitis. **d,** Intestinal weight:length for small intestine and colon were measured at terminal harvest of WT (n = 9) and Xist^TCD^ (n = 9). **e-f,** Representative H&E images **(left)** of and histologic severity scores (adapted from Erben et al) **(right)** for small intestine (n = 11) **(e)** and colon (n = 17) **(f).** Data are representative of 3 experiments. Samples were compared using Wilcoxon rank-sum test **(a)**, unpaired two-tailed Student’s *t-*test **(c-f)**. Error bars represent standard deviation. \*\**P* < 0.01; \*\*\**P* < 0.001.

To further explore Xist’s role in colitogenic T cell function, we monitored colitis development in RAG2 KO recipients, after separate transfer of WT or Xist^TCD^ CD45RB^hi^ CD4+ T cells (**Fig. 6B**). Recipients of WT donor cells developed robust colitis, as evidenced by significant weight loss **(Fig. 6C, gray)**. In contrast, recipients of Xist^TCD^ donor cells were protected from colitis, as evidenced by stable body mass compared to recipients of WT donor cells **(Fig. 6C, red)**. In accordance with preserved body mass, weight-to-length ratios of the small intestine and colon were significantly lower in recipients of Xist^TCD^ CD45RB^hi^ CD4+ T cells **(Fig. 6D)**, and histological sections of small intestine and colon demonstrated lower histological inflammation scores in recipients of Xist^TCD^ cells^52^, compared to WT cells **(Fig. 6E-F).** Finally, CD4+ T cell numbers were decreased in lymph nodes and spleens of recipients of Xist^TCD^ CD4+ T cells, consistent with a cell-intrinsic fitness defect **(Supp. Fig. 8D)**. Thus, Xist deficiency reduces the colitogenic capacity of CD45RB^hi^ CD4+ T cells. Importantly, B cells are absent in reconstituted hosts in this model. This is notable given previous work suggesting that Xist ribonucleoprotein complexes stimulate B cell autoantibody production to promote autoimmunity^14^. In contrast, our data indicate that cell-intrinsic Xist expression in CD4+ T cells promotes T cell-mediated colitis, even in the absence of B cells.

## DISCUSSION

While sex bias in autoimmune disease has been clearly demonstrated, the mechanisms driving increased autoimmune predisposition in females are not completely understood^53^. Here, we demonstrate a T cell-intrinsic role for the female-specific lncRNA, Xist, in promoting autoimmune predisposition. In T cells undergoing homeostatic expansion, T cell-specific Xist deletion derepresses X-linked immunoregulatory genes such as *Sh2d1a/*SAP and *Foxp3*. Moreover, T cell-specific Xist deletion increases susceptibility to activation-associated cell death, increases the frequency of Foxp3+ Tregs, and limits the colitogenic capacity of CD4+ T cells. Together, these findings support a model in which Xist-dependent regulation of X-linked gene expression helps sustain pathogenic CD4+ T cell responses.

Previous studies have reported non-canonical mechanisms by which Xist promotes autoimmunity. Beyond its role in repressing X-linked genes, Xist’s association with antigenic ribonucleoproteins (RNPs) has been reported to underlie its immunogenic function^14^. During SLE pathogenesis, neutrophils extrude Xist/RNP complexes, which serve as antigenic targets for autoantibodies^54^. Additionally, Xist has been proposed to function as a TLR7 agonist that stimulates type 1 IFN production by plasmacytoid dendritic cells^12^. Our findings implicate Xist’s canonical gene-silencing function in T cell regulation: T cell-specific Xist deletion increases *Sh2d1a*/SAP expression, including nascent transcription from the Xi, and reduces colitogenic activity.

The consequences of Xist-dependent silencing may differ across immune cell types. In B cells, XIST restrains *Tlr7* expression and the accumulation of atypical B cells^14^, whereas our findings suggest that Xist supports pathogenic T cell responses. In support of this, we demonstrate that CD4+ T cells from female patients with Crohn’s disease express higher levels of XIST than controls. This is in line with previous publications, which report that other female-biased autoimmune diseases are also associated with higher XIST levels in peripheral blood mononuclear cells^12,13^. In particular, higher XIST levels were noted in NK cells and CD4+ T cells of primary biliary cholangitis patients^13^ and in B cells of SLE patients^12^. Thus, the effects of Xist on immune function are likely to depend on both the cellular context and the genes whose expression it controls, alongside its immunogenic functions.

We found that T cell-specific Xist deletion results in upregulation of *Sh2d1a*, which encodes the X-linked immunoregulatory protein SAP. As an SH2 adaptor that binds SLAM family receptors, SAP plays a key role in T cell signaling, activation, and subsequent apoptosis-mediated T cell contraction^42,55^. T cell contraction plays a critical role in constraining effector T cell responses after activation. Left unchecked, the effector T cell response can lead to serious and even fatal tissue damage. Notably, SAP is deficient in X-linked Lymphoproliferative Disease (XLD), a condition characterized by over-expansion of lymphocytes that is triggered by viral infection in childhood^56^. The importance of SAP in controlling the immune response is highlighted by the high rate of lethality in SAP-deficient individuals, which is often associated with lymphadenopathy, hepatosplenomegaly, and tissue infiltration^56^. Whether increased SAP expression is required for the loss of T cell fitness after Xist deletion remains unresolved. Testing whether normalization of SAP expression restores fitness and colitogenic activity would establish its contribution to these phenotypes.

Previous work showed that Xist is required for maintenance of X-linked gene silencing in B cells, and that X-linked targets of Xist are enriched for ‘escapees’^8^. Despite chromosome-wide silencing with XCI, these escapees continue to be expressed from the Xi and are estimated to represent about 15% of X-linked genes in humans and 3% of X-linked genes in mice^57^. In mice under steady-state conditions, escapees are similarly over-represented among Xist-dependent genes in CD4+ T cells. In contrast, a smaller proportion of Xist-dependent genes are escapees in CD4+ T cells after their expansion in lymphopenic hosts. *Sh2d1a* and *Foxp3* for instance, have not previously been shown to be expressed from the Xi, and are therefore not known escape genes^10^. These findings suggest that the repertoire of genes dependent on Xist for silencing varies with T cell state. How T cell activation state alters the identity of genes requiring Xist-dependent gene silencing, however, remains unclear and will require further study.

In our study, we found upregulation of *Foxp3* transcripts in CD4+ T cells at steady state and undergoing homeostatic proliferation. In the setting of colitis, however, Foxp3 transcripts were not increased despite a significant increase in the frequency of Foxp3+ CD4+ T cells by flow cytometry. This discrepancy may reflect the low overall frequency of Tregs among total CD4+ cells. Despite their low frequency, induced Tregs are an essential protective cell type in autoimmune colitis as their ablation results in exacerbated disease ^58^. Further investigation is needed to explore the role of Xist-driven regulation of FOXP3 in the modulation of autoimmune severity.

Finally, our findings may contribute to efforts to develop sex-informed biomarkers of autoimmune disease. It has been proposed that anti-XIST-associated autoantibodies (AXA) can be useful as a diagnostic and prognostic markers for autoimmune disease in females^54^, based on the finding that extrusion of XIST RNP particles by neutrophils in female patients with autoimmune disease leads to the production of AXAs. Adding to this, we propose that Xist levels may represent another potential biomarker, given our data that Xist levels are increased in CD4+ T cells of patients with Crohn’s disease. These data build upon previous data from other groups reporting elevated Xist levels in peripheral blood mononuclear cells of autoimmune disease patients^12,13^. Together, these findings may complement recent efforts to develop “XIST-aware diagnostics” for the benefit of patients with autoimmunity.

## FIGURES

**Supp. 1:**
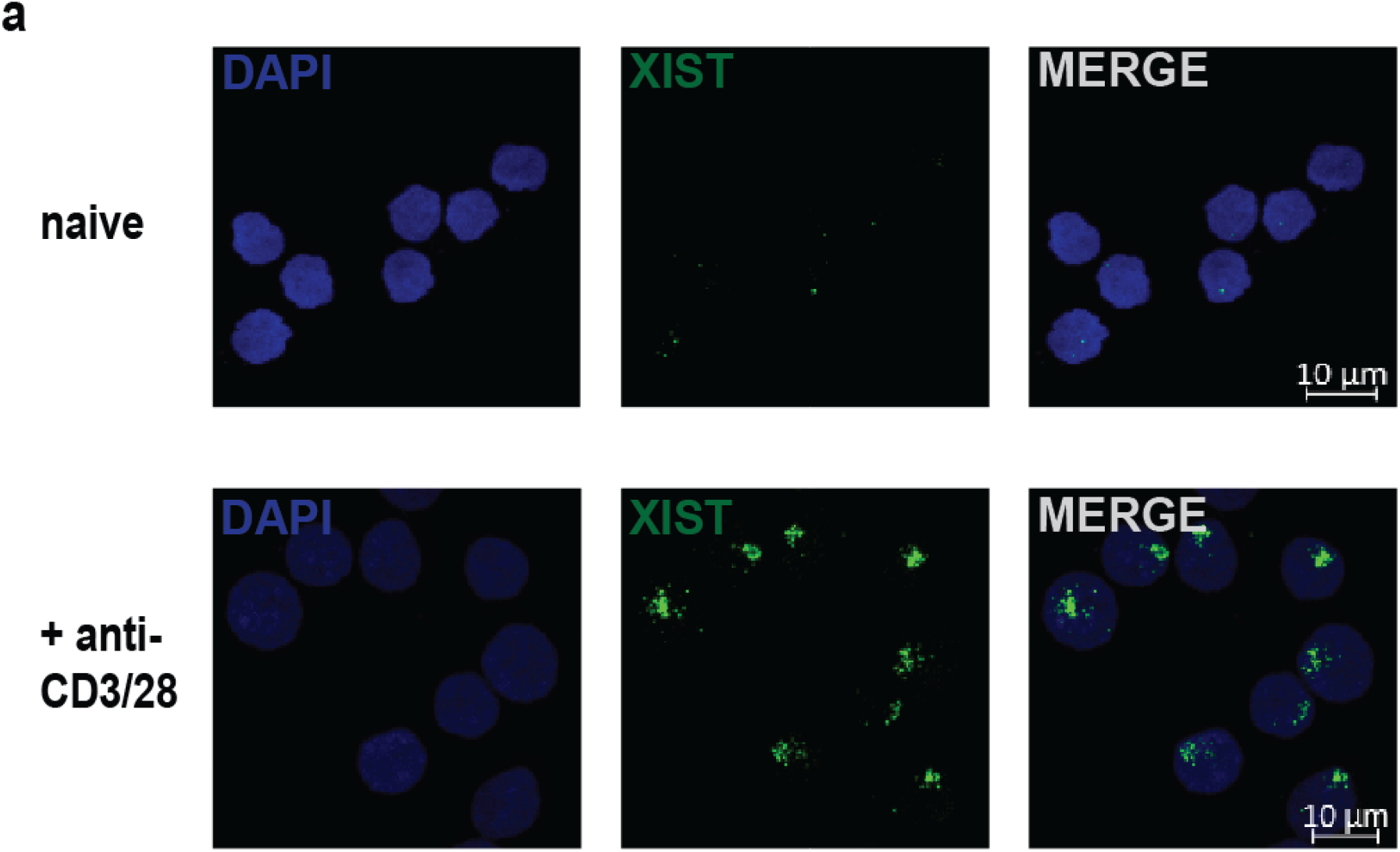
Xist localization in human CD4+ T cells is state dependent. **a**, Xist fluorescent in situ hybridization (FISH) in naïve **(top)** and activated (48 hours, 2 ug/mL anti-CD3/28) **(bottom)** human CD4+ T cells.

**Supp. 2:**
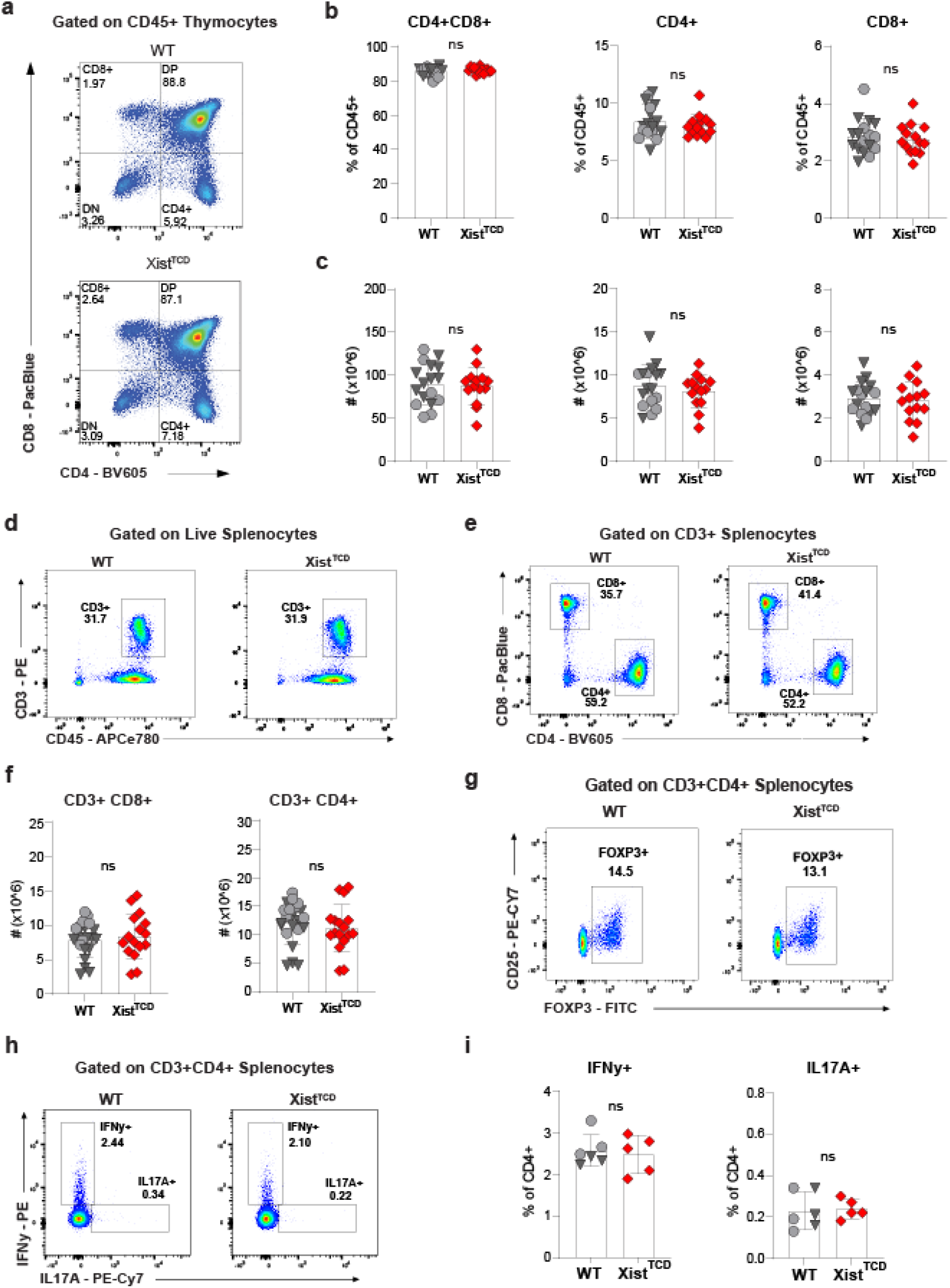
Xist minimally affects thymic T cells and splenic T helper subsets in steady-state. **a-c,** Flow cytometry of WT (n = 17) and Xist^TCD^ (n = 14) thymocytes from 8-week-old animals. Representative dot plots **(a),** percentage **(b),** and absolute number **(c)** of thymic subsets**. d-i,** Flow cytometry of WT and Xist^TCD^ splenocytes from 8-week-old animals. Representative dot plots showing percentage CD3+ T cells of live splenocytes **(d)** and percentage CD8+ or CD4+ of CD3+ T cells **(e). f,** Absolute number of CD3+CD8+ and CD3+CD4+ splenocytes in WT (n = 17) and Xist^TCD^ (n = 14) animals. **g,** Representative dot plots showing percentage Foxp3+ of CD3+CD4+ splenocytes. **h-i,** Representative dot plots **(h)** and percentage IFNy and IL17A of WT (n = 6) and Xist^TCD^ (n = 5) CD3+CD4+ splenocytes **(i)**. Data are representative of 2-4 experiments. Samples were compared using unpaired two-tailed Student’s *t-*test **(b-c; f; i)**. Error bars represent standard deviation.

**Supp. 3:**
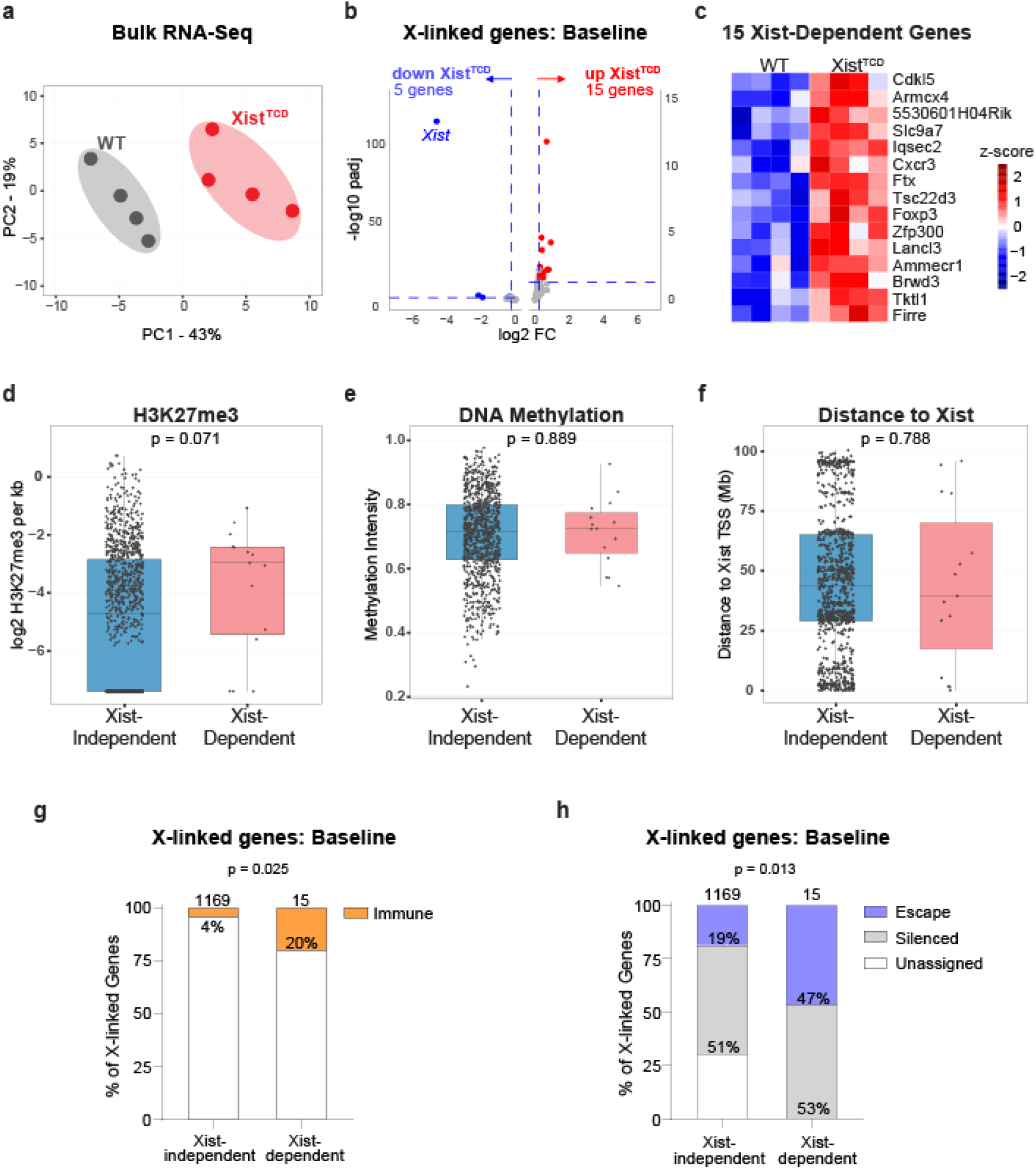
T cell Xist silences X-linked ‘escapees’ in steady-state. Bulk RNA-Seq was performed on splenic CD4+ T cells from 8-week WT and Xist^TCD^ animals (n = 4, samples are biologic replicates). Principal component analysis **(a)**, volcano plot of X-linked genes **(b)**, and heatmap of Xist-dependent genes (X-linked, padj < 0.05, log2FC > 0.25) **(c)** in steady-state WT and Xist^TCD^ CD4+ T cells. **d-f,** H3K27me3 **(d),** DNA methylation **(e),** and X chromosome location (distance to Xist TSS) **(f)** for Xist-dependent genes versus Xist-independent genes (remaining X-linked genes). Percentage immune genes **(g)** and escape or silenced genes **(h)** among Xist-dependent genes versus Xist-independent. Comparisons were made using two-sided Fisher’s exact test **(d)**, Wilcoxon rank-sum test **(e-f)**, and one-sided Fisher’s exact test **(g-h)**.

**Supp. 4:**
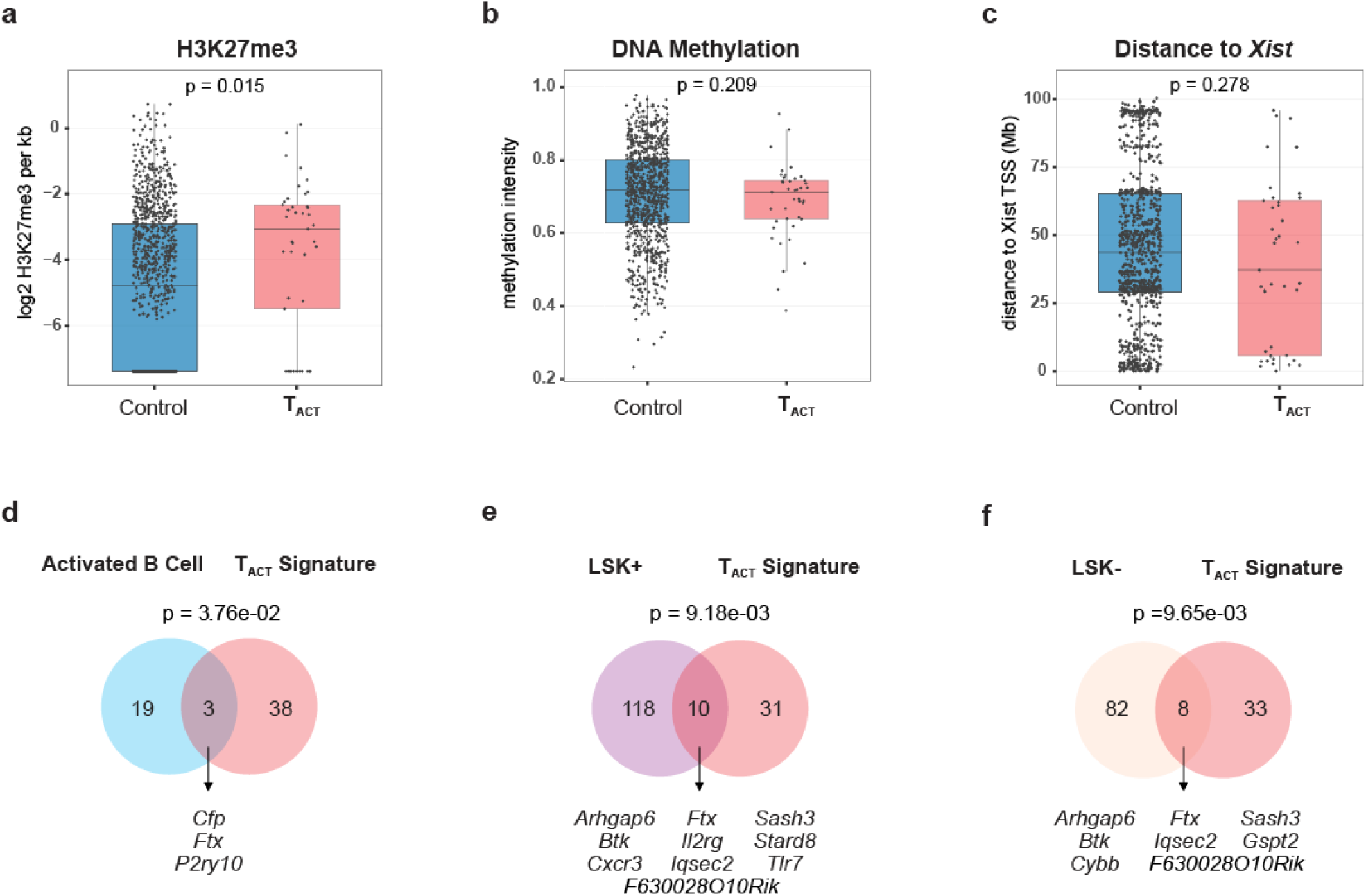
Xist targets a distinct set of genes in activated T cells. Analysis of T_Act_ signature genes (X-linked genes, p < 0.05, log2FC > 0.25, common to 6 and 12 week CD3+ T cell adoptive transfer datasets) **a-c,** H3K27me3 **(a),** DNA methylation **(b),** and X chromosome location (distance to Xist TSS) **(c)** for T_Act_ signature genes versus control (remaining X-linked genes)**. d-f,** Overlap of T_Act_ signature genes with Xist-dependent genes in activated B cells **(d)** and hematopoietic stem cells [LSK+ (Lin- [LSK+ (Lin-, Sca-1+, c-Kit+) and LSK- (Lin-, Sca-1-, c-Kit+)] **(e-f)**. Comparisons were performed using two-sided Fisher’s exact test **(a)**, Wilcoxon rank-sum test **(b-c)**, and one-sided Fisher’s exact test **(d-f)**.

**Supp. 5:**
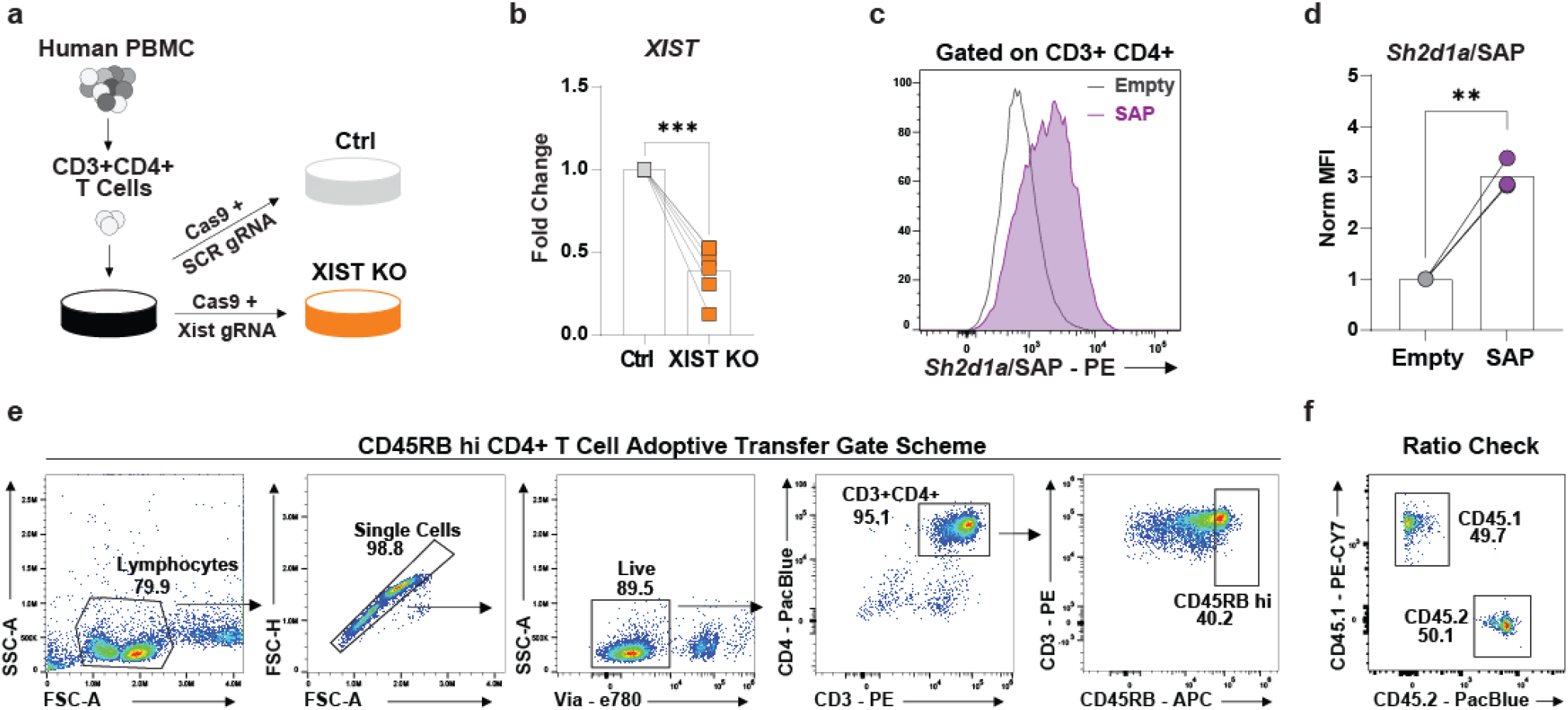
Validation of XIST CRISPR editing, *Sh2d1a*/SAP overexpression, and CD45RB hi CD4+ T cell isolation. **a,** Schematic of CRISPR editing workflow**. b,** *XIST* expression by qPCR in Ctrl or XIST KO human CD4+ T cells one week after electroporation. **c,** Representative *Sh2d1a*/SAP histogram of control-transduced (MSCV-EMPTY-IRES-mCherry) and SAP-overexpressing (MSCV-Sh2d1a-IRES-mCherry) CD3+CD4+ T cells. **d,** Sh2d1a/SAP MFI normalized to control-transduced CD3+CD4+ T cells (n = 3). **e,** Flow sorting gate scheme for isolation of CD45RB hi CD4+ T cells for adoptive transfer colitis. **f**. Representative flow plot showing the relative frequency of congenically-marked CD45RB hi CD4+ donor T cell populations. The ratio of donor populations was checked before each co-transfer experiment. Samples were compared using paired two-tailed Student’s *t-*test **(b,d)**. \*\**P* < 0.01 \*\*\**P* < 0.001.

**Supp. 6:**
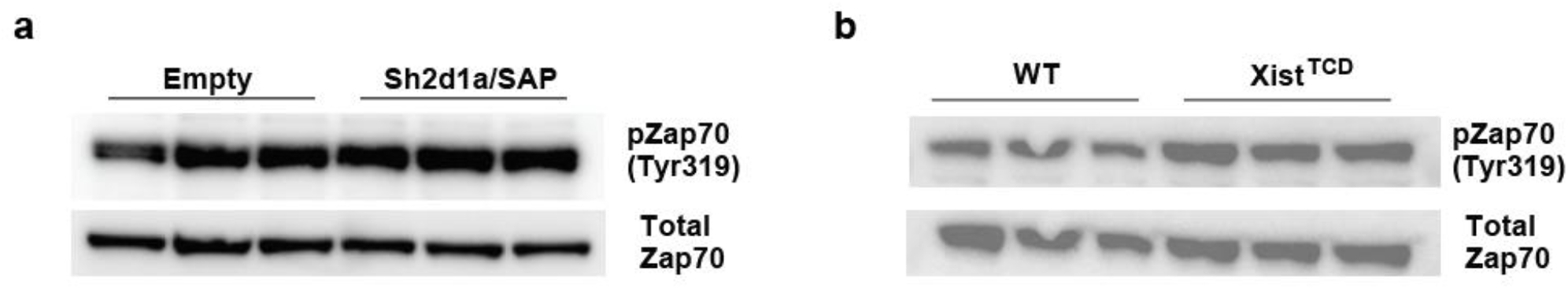
Xist modulates T cell receptor signaling in CD4+ T cells. Western blot for pZap70 (Tyr319) and total Zap70 in control-transduced (MSCV-Empty-IRES-mCherry) versus Sh2d1a/SAP-overexpressing (MSCV-Sh2d1a-IRES-mCherry) **(a)** or WT versus Xist^TCD^ **(b)** CD4+ T cells ninety minutes after plate bound anti-CD3/28 antibody restimulation.

**Supp. 7:**
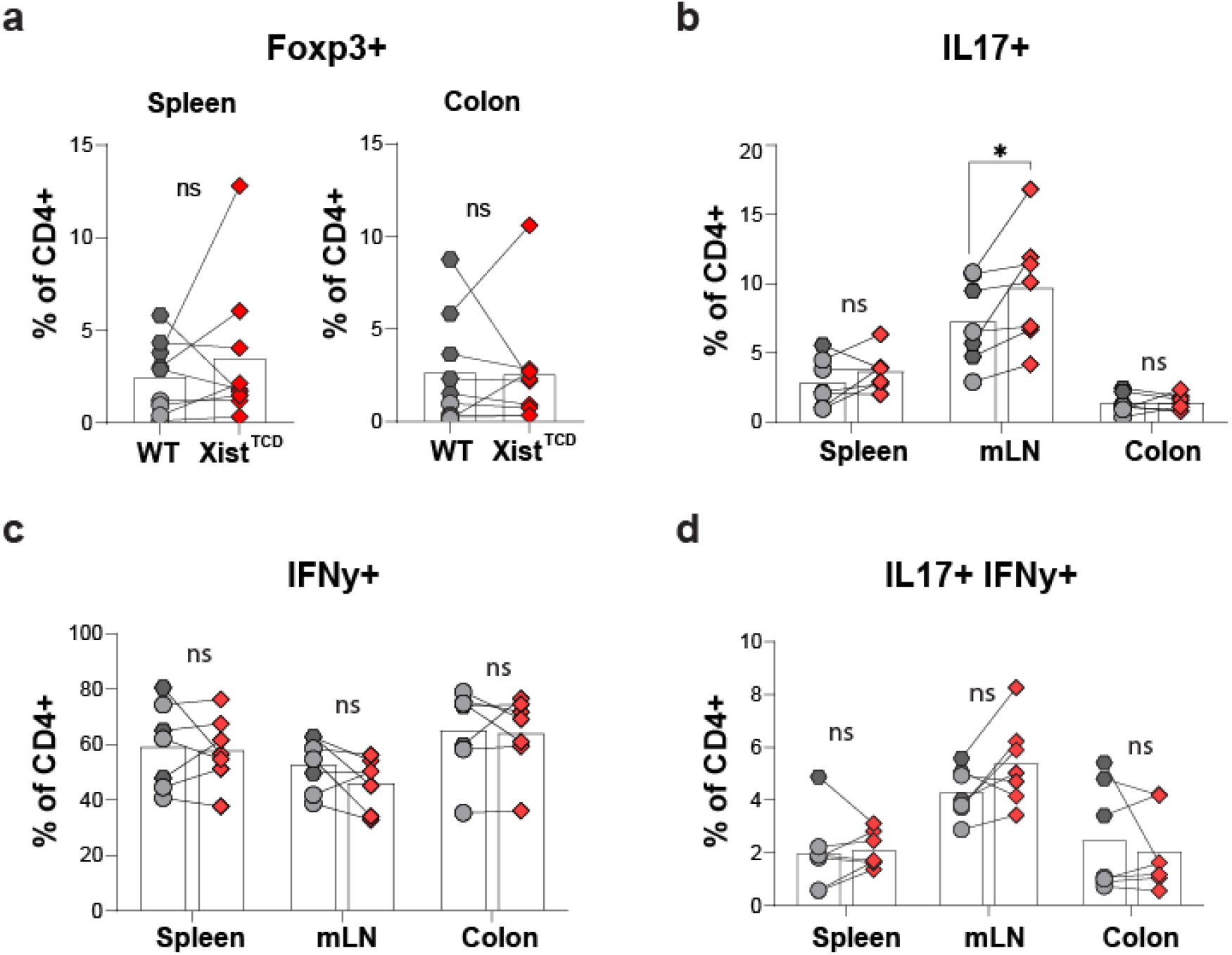
Xist minimally affects CD4+ T cell cytokine production in autoimmune colitis. Analysis of WT and Xist^TCD^ CD4+ T cells across tissues in RAG2 KO recipients 7 weeks after co-transfer of WT and Xist^TCD^ CD45RB hi CD4+ T cells. Percentage Foxp3+ of WT (n = 9) and Xist^TCD^ (n = 9) CD4+ T cells in the spleen and colon **(a)**. Frequency IL-17-producing **(b),** IFNy-producing **(c)**, and IL-17/IFN-y-coproducing **(d)** WT (n = 7) and Xist^TCD^ (n = 7) CD4+ T cells in RAG2 KO recipient spleen, mLN, and colon. Data are representative of two experiments. Comparisons were performed using paired two-tailed Student’s *t-*test **(a-d)**. \**P* < 0.05.

**Supp. 8:**
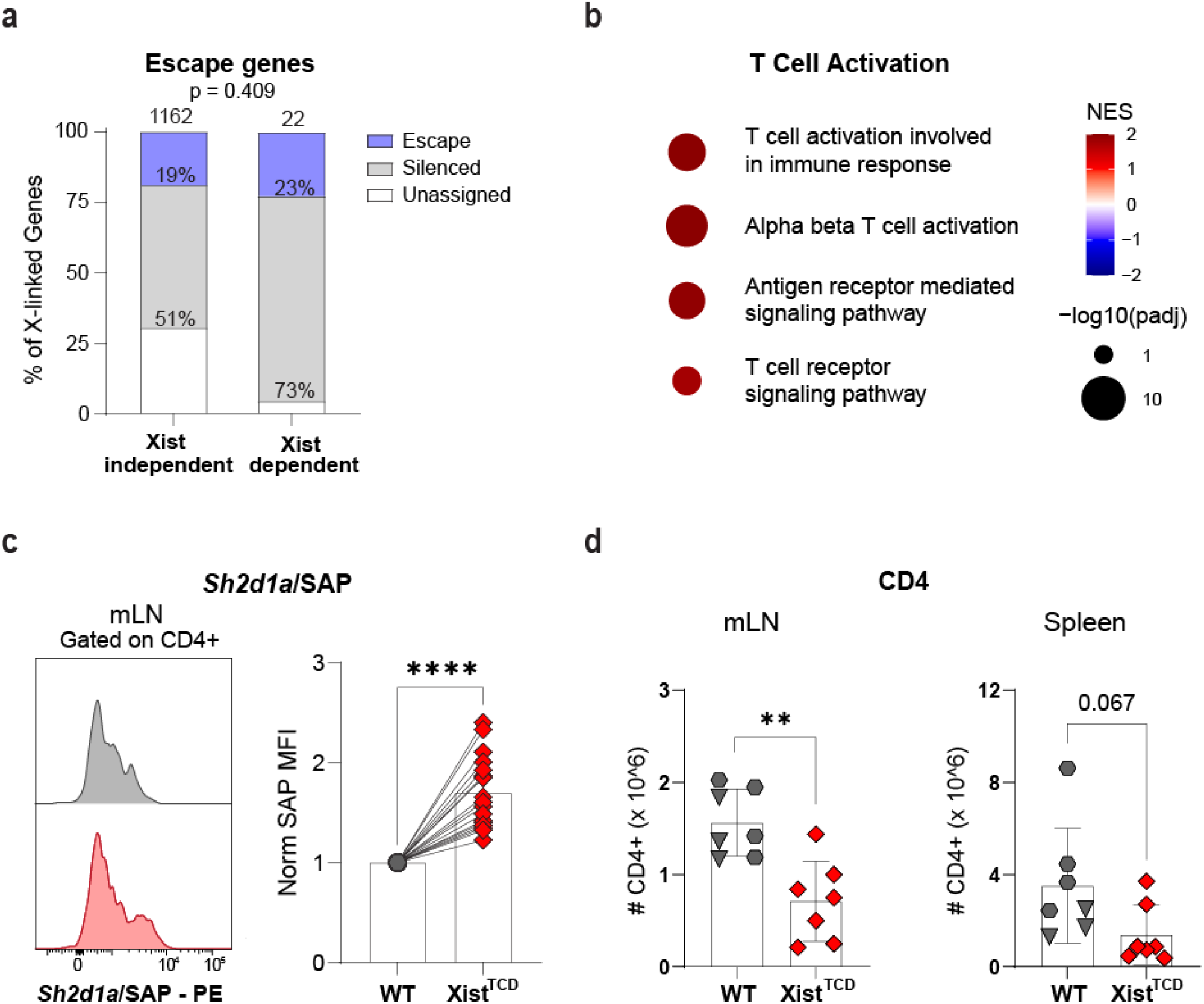
T Cell Xist regulates Sh2d1a/SAP expression and colitogenic CD4+ T cell fitness. **a,** Proportion of escape and silenced genes among Xist-dependent genes (X-linked genes, p < 0.05, log2FC > 0.25) common to colon and mLN and Xist-independent genes (remaining X-linked genes). **b,** Gene set enrichment analysis (GSEA) of WT and Xist^TCD^ CD4+ T cells from the colon of RAG2 KO recipients 7 weeks after CD45RB hi CD4+ T cell co-transfer [1:9 WT (CD45.1) and Xist^TCD^ (CD45.2), to ensure adequate numbers of Xist-deficient cells]. Dot plot of select biological processes gene sets. **c,** Representative Sh2d1a/SAP histograms and normalized Sh2d1a/SAP MFI in WT and Xist^TCD^ CD4+ T cells from the mLN of RAG2 KO animals (n = 17) reconstituted with WT and Xist^TCD^ CD45RBhi CD4+ T cells. **d,** CD4+ T cell numbers in mesenteric lymph node and spleen of RAG2 KO recipients (n = 7) 9 weeks after adoptive transfer of WT or Xist^TCD^ CD45RB hi CD4+ T cells. Data are representative of 2-4 experiments. Comparisons were performed using Fisher’s exact test **(a)**, paired two-tailed Student’s *t-*test **(c)**, and unpaired two-tailed Student’s *t-*test **(d)**. Error bars represent standard deviation. \*\**P* < 0.01; \*\*\*\**P* < 0.0001

## MATERIALS AND METHODS

### Mice

Mice were maintained at UCLA following Institutional Animal Care and Use Committee (IACUC) guidelines and Animal Research Committee (ARC)-approved protocols (ARC-2018-082, ARC-2026-027, ARC-2025-018). The following mouse strains were utilized: B6 CD45.2 (C57BL/6J; Jackson Labs: 000664), B6 CD45.1 (B6.SJL - Ptprca Pepcb/BoyJ; Jackson Labs: 002014), 129 Xist 2lox (129-*Xist^tm2Jae^*/Mmnc, Mutant Mouse Resource & Research Centers: 029172-UNC), CD4 Cre [B6.Cg-Tg(Cd4-cre)1Cwi/BfluJ, Jackson Labs: 022071], RAG2 KO (B6.Cg-Rag2tm1.1Cgn/J, Jackson Labs: 008449), and Nur77-GFP [C57BL/6-Tg(Nr4a1-EGFP/cre)820Khog/J, Jackson Labs: 016617]. 129 Xist 2lox mice were backcrossed to B6 CD45.2 mice for at least 6 generations. B6 Xist 2lox mice were crossed with CD4 Cre mice to generate Xist^T^ ^cell^ ^deficient^ ^(TCD)^ mice (*CD4* ^Cre/WT^ *Xist* ^FL/FL^). B6 CD45.1x2 congenic mice were generated de novo by crossing of B6 CD45.2 and B6 CD45.1. 6- to 12-week age-matched female mice were used for experiments. Littermate Cre negative WT controls were used when possible and experimental cohorts were randomly assigned.

### Isolation and adoptive transfer of mouse T cells

Mouse spleens were collected and processed to single-cell suspensions as previously described. Red blood cell lysis was performed prior to T cell isolation. CD3+ (Stemcell Technologies, 19851) or CD4+ (Stemcell Technologies, 19852) T cells were isolated via immunomagnetic negative selection per the manufacturer’s protocol. For CD3+ T cell co-adoptive transfer experiments, 1x10^6 CD3+ T cells of each genotype were transferred to RAG2 KO recipients via retro-orbital injection. Flow cytometry was performed to measure the purity and relative ratio of donor cells prior to injection.

### Flow cytometry and fluorescence activated cell sorting

Fluorophore-conjugated antibodies were used for analysis of cell surface, intracellular, and intranuclear proteins. Cell surface marker staining was performed in FACS buffer (2% FBS and 2mM EDTA in PBS). Intracellular and intranuclear staining were performed by fixing and permeabilizing cells with the Cytofix/Cytoperm kit (BD, 554714) or Transcription Factor kit (eBioscience, 00-5523-00), respectively. Annexin V staining was performed using Annexin V binding buffer (Biolegend, 422201) and all samples were analyzed within 1 hour of staining completion. BD LSR Fortessa was used for flow cytometry and BD FACS Aria or Cytek Aurora CS for cell sorting. Data analysis was performed using FlowJo software.

### Bulk RNA-seq

RNA was extracted from immunomagnetic- and FACS-isolated T cells using Quick-RNA Microprep Kit (Zymo Research, R1051). RNA quality was assessed using High Sensitivity RNA Screen Tape (Agilent). RNA-seq library preparation was performed using ABclonal FAST RNA-seq Lib Prep Kit. 2x50 paired-end sequencing was performed using the NovaSeq X Plus Sequencing System.

Paired-end reads were aligned to the mouse genome (mm10) using HISAT2 and raw counts were obtained using featureCounts. Raw counts were utilized for differential expression analysis by DESeq2. For GSEA analysis, the msigdbr package was used to retrieve the Hallmark and GO Biological Processes Molecular Signatures Database (MSigDB) gene sets. The fgsea function was run with minSize and maxSize of 15 and 1000, respectively.

### Colitis induction

Colitis was induced in RAG2 KO mice by adoptive transfer of 5 x 10^5 CD4+ CD45RB hi donor cells as previously described^59^. In brief, splenocyte single-cell suspensions from donor mice were subjected to immunomagnetic negative selection (Stemcell Technologies, 19852) to enrich for CD3+ CD4+ T cells. The top 40% of CD45RB-expressing CD3+CD4+ T cells were then sorted by FACS and administered to recipient animals by intraperitoneal injection. Recipient mouse body weight was measured bi-weekly to track disease progression. Animals with a 20% reduction from initial body weight were sacrificed.

### Intestinal tissue processing and histology

Small intestine and large intestine were collected for downstream analysis. For flow cytometry and cell sorting lamina propria-infiltrating immune cells were isolated as previously described. In short, intestinal samples were washed in DTT solution [5mMEDTA, 1mM DTT, 10% FCS, 15mM HEPES in RPMI (37C, 225rpm, 45 minutes)], digested in collagenase solution [200U/mL collagenase type 1A (SIGMA), 10% FCS, 15 mM HEPES in RPMI (37C, 225 rpm, 30 minutes)], and subjected to density centrifugation in percoll solution [40% osmotically-adjusted percoll in PBS (22C, 600xG, 20 minutes)].

For histologic analysis, small and large intestines were prepared using the swiss roll technique with formalin-fixation and paraffin-embedding. Routine H&E-stained sections were scored for disease activity with a previously published histomorphological scale^60^ based on six criteria [severity and extent of inflammatory cell infiltrate, crypt expansion, goblet cell loss, crypt distortion, and villous blunting (small intestine only)] to generate a cumulative severity score ranging from 0-21 for large intestine and 0-26 for small intestine. Scoring was performed by a practicing gastrointestinal pathologist (DD) blinded to experimental cohorts.

### Cloning, retrovirus production, and transduction

Retroviral vectors for Sh2d1a overexpression were cloned using Gibson Assembly (NEBuilder HiFi DNA Assembly). Sh2d1a was PCR amplified (GenScript, Omu18688) and inserted into restriction enzyme-digested retroviral backbones [MSCV-IRES-GFP (Addgene #20672) & MSCV-IRES-mCherry (Addgene #52114). Final plasmid sequences were validated by sequencing (Plasmidsaurus).

For retroviral production, cloned sap-overexpressing (MSCV-Sh2d1a-IRES-GFP, MSCV-Sh2d1a-IRES-mCherry) or control plasmids (MSCV-IRES-GFP, MSCV-IRES-mCherry) were added to HEK293T cells with a retroviral packaging plasmid (pCL-ECO, Addgene #12371) and TransIT-293 transfection reagent (Mirus Bio). The day after transfection, sodium butyrate was added to HEK293T cells to enhance retroviral output. Two days later, viral supernatants were collected and titered as previously described.

For T cell transduction, spinoculation was performed. Viral supernatant and activated T cells were added to RetroNectin-coated plates (Takara Bio) and centrifuged (2000G, 32C, 90 minutes). Following spinoculation, T cells were cultured on anti-CD3/28 (Invitrogen, 14-0031 and 14-0281, 1 ug/mL each) coated plates until downstream experimental use.

### T cell restimulation assays

CD3+ CD4+ T cells were initially activated for 3-5 days on anti-CD3/28 (Invitrogen, 14-0031 and 14-0281, 1 ug/mL each) coated plates in complete RPMI and subsequently rested on uncoated plates in complete RPMI supplemented with hIL2 (50 U/mL). Restimulation was performed with soluble anti-CD3 for flow cytometry and plate-bound anti-CD3/28 for western blot.

### Western blot

Protein was extracted from cell pellets using Pierce RIPA buffer (Thermo Fisher) with Halt protease and phosphatase inhibitor cocktail (Thermo Fisher) and quantified using the Pierce BCA Protein Assay Kit (Thermo Fisher). Samples were electrophoresed on NuPage Novex 4–12% Bis-Tris Protein Gels, transferred to PVDF membranes, and blocked overnight with 5% wt/vol milk in 1x TBS and 0.1% Tween-20. Immunoblots were performed using rabbit anti-pZap70 (Tyr319) (1:1000; Cell Signaling, 2701), rabbit anti-Zap70 (1:1000; Cell Signaling, 2705) and goat anti-rabbit IgG HRP secondary antibody (1:10,000; Thermo Fisher, 31466). SuperSignal West Pico PLUS ECL (Thermo Fisher) was used for protein detection and the Azure Biosystems c280 system was used for imaging.

### Single cell RNA-seq

For analysis of intestinal CD4+ T cells from Crohn’s disease patients and healthy controls, a previously published dataset was used. A Seurat object was generated from terminal ileum and small bowel immune cells. Filtering excluded cells with > 200 features, < 4000 features, > 300 counts, and < 25% mitochondrial reads. The filtered object was subset to include only CD4+ T cells (using anno2 cluster identities established by Kong et al) from female donors. The female CD4+ T cell object was subject to normalization, variable feature identification (nfeatures = 2,000), scaling (regression of “percent.mt” and “nCount_RNA”), and principal component analysis (npcs = 30) following the standard Seurat pipeline. For analysis of Xist expression, cells with XIST > 0 were evaluated.

### RNA FISH

RNA FISH was performed as previously described ^27^. Probes for RNA FISH experiments were labeled using the BioPrime DNA Labeling System (Invitrogen, cat. no. 18094011), according to the manufacturer’s instructions. Briefly, 100-1,000 ng of DNA was labeled in a 40 µL reaction containing 1x random primers, heated to 95°C for 5 mins and then snap cooled on ice. To the reaction, we added 5 µL fluorescently labeled dUTPs [Thermo Fisher, cat. no. C11399 (568-5-dUTP) and C11397 (488-5-dUTP), and GE Healthcare, cat. no. PA55022 Cy5-dUTP)], 5 µL AGC ^1^/_2_T dNTPs, and 2 µL of Klenow exo-enzyme, and incubated in the dark at 37°C overnight. The reaction was purified using AMPure beads. The probes were then resuspended in nuclease-free water and ethanol-precipitated with yeast tRNA, and sodium acetate at -80°C overnight. After washing with 100% ethanol, probes were resuspended in nuclease-free water and ethanol-precipitated with unlabeled sheared salmon sperm DNA (Sigma, cat no. D9156), Cot1 DNA (Invitrogen, cat. no. 1844016), and yeast tRNA at -80°C, overnight. After an ethanol series (70%–100%), probes were resuspended in deionized formamide with shaking at 37°C for 6 hours. Probes were then adjusted in hybridization buffer (50% formamide, 2 × SSC, 10% dextran sulfate) and stored at −80°C.

Labeled DNA probes for *Sh2d1a* were generated from four PCR-amplified segments of intronic regions, each approximately 5 kb in length. The primer pairs used for these PCR reactions were as follows: segment 1, F: GCTCTGTTTGCGTTGCTGGCAC, R: TGAGCCAATCCAGGGAAGTGGC; segment 2, F: GCCACTTCCCTGGATTGGCTCA, R: TCCTGTGCCTAAGAACAACAGA; segment 3, F: TCCTGTGCCTAAGAACAACAGA, R: GCACAGAGTCCCACCTCTAGCCA; and segment 4, F: TGGCTAGAGGTGGGACTCTGTGC, R: TGAGCCCTGAAGCAACTTGGCA. Labeled probes for *Atrx* were generated from BAC DNA (RP23-265D6). Labeled probes for mouse Xist and human XIST were generated from the respective full-length cDNAs. All cells were counterstained with DAPI. Images were acquired on a Zeiss LSM 880 confocal microscope. ImageJ was used for processing z-stack images.

### Allelic classification

RNA FISH for *Atrx* and Xist was used to identify whether *Sh2d1a* nascent transcripts arose from the Xa or Xi. In the WT CD4+ T cells, cells with the *Sh2d1a* nascent transcript signal colocalizing with a single *Atrx* transcription site on the Xist-negative chromosome were classified as exhibiting Xa-only expression, whereas cells with the nascent *Sh2d1a* signal colocalizing exclusively with Xist RNA were classified as exhibiting Xi-only expression. Cells with *Sh2d1a* transcription signals associated with both chromosomes were classified as exhibiting biallelic expression. In Xist-deficient cells (Xist^TCD^ cells), in which Xist RNA is absent, cells were considered to have Xi expression if the *Sh2d1a* signal did not colocalize with the single *Atrx* transcription signal. Cells where *Atrx* exhibited biallelic expression (16% in WT cells and 17% in Xist^TCD^ cells) or Xi expression (2% in WT cells) were excluded from the analysis.

### CRISPR

Human CD4+ T cells were isolated from PBMCs via immunomagnetic isolation (Stemcell, 17952) and stimulated with plate-bound anti-CD3/28 (1ug/mL each) overnight in preparation for CRISPR editing. Ribonucleoprotein (RNP) complexes were prepared just prior to transfection. Specifically, Xist-targeting^8^ or scramble guide RNA (gRNA) and recombinant nuclear-localized sequence-tagged Cas9 (NLS-Cas9) protein (QB3 MacroLab) were mixed at a 2.5:1 molar ratio and incubated at 37C for 15 minutes. CD4+ T cells were harvested from culture, washed, resuspended in electroporation buffer, and mixed with the pre-assembled RNP complexes. CD4+ T cells were electroporated using the Neon NxT Electroporation System (1400 volts, 10 ms, 3 pulses). Immediately following electroporation, CD4+ T cells were added to pre-warmed complete RPMI and rested at 37C for 30 minutes. Subsequently, cells were added to anti-CD3/28 (1ug/mL each) coated plates for the start of the T cell restimulation assay.

### Statistics

Statistical analysis was performed using GraphPad Prism. Statistical tests for each experiment are stated in figure legends. For bulk RNA-seq, per-gene two-sided Wald test with Benjamini-Hochberg FDR correction was performed to adjust for multiple comparisons. For single cell RNA-seq, Wilcoxon rank-sum test was utilized for comparison of XIST expression. Sample sizes are reported in figure legends. Unless otherwise specified, samples represent single measurements from biologic replicates.

### Data and Code Availability

Raw bulk RNA-seq data are available on the NCBI Gene Expression Omnibus (GSE350340).

## REFERENCES

1 Fairweather, D., Frisancho-Kiss, S. & Rose, N. R. Sex differences in autoimmune disease from a pathological perspective. Am J Pathol 173, 600–609 (2008). 10.2353/ajpath.2008.071008

2 Brockdorff, N., Bowness, J. S. & Wei, G. Progress toward understanding chromosome silencing by Xist RNA. Genes Dev 34, 733–744 (2020). 10.1101/gad.337196.120

3 Jegu, T., Aeby, E. & Lee, J. T. The X chromosome in space. Nat Rev Genet 18, 377–389 (2017). 10.1038/nrg.2017.17

4 Loda, A., Collombet, S. & Heard, E. Gene regulation in time and space during X-chromosome inactivation. Nat Rev Mol Cell Biol 23, 231–249 (2022). 10.1038/s41580-021-00438-7

5 Plath, K., Mlynarczyk-Evans, S., Nusinow, D. A. & Panning, B. Xist RNA and the mechanism of X chromosome inactivation. Annu Rev Genet 36, 233–278 (2002). 10.1146/annurev.genet.36.042902.092433

6 Robert Finestra, T. & Gribnau, J. X chromosome inactivation: silencing, topology and reactivation. Curr Opin Cell Biol 46, 54–61 (2017). 10.1016/j.ceb.2017.01.007

7 Brown, C. J. & Willard, H. F. The human X-inactivation centre is not required for maintenance of X-chromosome inactivation. Nature 368, 154–156 (1994). 10.1038/368154a0

8 Yu, B. et al. B cell-specific XIST complex enforces X-inactivation and restrains atypical B cells. Cell 184, 1790–1803 e1717 (2021). 10.1016/j.cell.2021.02.015

9 Yang, L., Yildirim, E., Kirby, J. E., Press, W. & Lee, J. T. Widespread organ tolerance to Xist loss and X reactivation except under chronic stress in the gut. Proc Natl Acad Sci U S A 117, 4262-4272 (2020). 10.1073/pnas.1917203117

10 Hauth, A. et al. Escape from X inactivation is directly modulated by Xist noncoding RNA. Nat Cell Biol 28, 166–181 (2026). 10.1038/s41556-025-01823-6

11 Huret, C. et al. Altered X-chromosome inactivation predisposes to autoimmunity. Science Advances 10, eadn6537 (2024). doi:10.1126/sciadv.adn6537

12 Crawford, J. D. et al. The XIST lncRNA is a sex-specific reservoir of TLR7 ligands in SLE. JCI Insight 8 (2023). 10.1172/jci.insight.169344

13 She, C. et al. Effect of LncRNA XIST on Immune Cells of Primary Biliary Cholangitis. Front Immunol 13, 816433 (2022). 10.3389/fimmu.2022.816433

14 Dou, D. R. et al. Xist ribonucleoproteins promote female sex-biased autoimmunity. Cell 187, 733–749 e716 (2024). 10.1016/j.cell.2023.12.037

15 Suárez-Fueyo, A., Bradley, S. J. & Tsokos, G. C. T cells in Systemic Lupus Erythematosus. Curr Opin Immunol 43, 32–38 (2016). 10.1016/j.coi.2016.09.001

16 Nakagawa, R. et al. CD4(+) T cells from patients with primary biliary cholangitis show T cell activation and differentially expressed T-cell receptor repertoires. Hepatol Res 49, 653–662 (2019). 10.1111/hepr.13318

17 Chan, J. E. et al. Shared CD4+ T cell receptor specificity groups in Crohn’s disease and ulcerative colitis. JCI Insight 11 (2026). 10.1172/jci.insight.195354

18 Syrett, C. M. et al. Altered X-chromosome inactivation in T cells may promote sex-biased autoimmune diseases. JCI Insight 4 (2019). 10.1172/jci.insight.126751

19 Wang, J. et al. Unusual maintenance of X chromosome inactivation predisposes female lymphocytes for increased expression from the inactive X. Proc Natl Acad Sci U S A 113, E2029–2038 (2016). 10.1073/pnas.1520113113

20 Jiwrajka, N. et al. Impaired dynamic X-chromosome inactivation maintenance in T cells is a feature of spontaneous murine SLE that is exacerbated in female-biased models. J Autoimmun 139, 103084 (2023). 10.1016/j.jaut.2023.103084

21 Booth, C. et al. X-linked lymphoproliferative disease due to SAP/SH2D1A deficiency: a multicenter study on the manifestations, management and outcome of the disease. Blood 117, 53–62 (2011). 10.1182/blood-2010-06-284935

22 Uhlig, H. H. et al. The diagnostic approach to monogenic very early onset inflammatory bowel disease. Gastroenterology 147, 990–1007 e1003 (2014). 10.1053/j.gastro.2014.07.023

23 Patey-Mariaud de Serre, N., et al. Digestive histopathological presentation of IPEX syndrome. Mod Pathol 22, 95–102 (2009). 10.1038/modpathol.2008.161

24 Gomez-Bris, R. et al. CD4 T-Cell Subsets and the Pathophysiology of Inflammatory Bowel Disease. Int J Mol Sci 24 (2023). 10.3390/ijms24032696

25 Syrett, C. M. et al. Diversity of Epigenetic Features of the Inactive X-Chromosome in NK Cells, Dendritic Cells, and Macrophages. Front Immunol 9, 3087 (2018). 10.3389/fimmu.2018.03087

26 Lee, P. P. et al. A critical role for Dnmt1 and DNA methylation in T cell development, function, and survival. Immunity 15, 763–774 (2001). 10.1016/s1074-7613(01)00227-8

27 Markaki, Y. et al. Xist nucleates local protein gradients to propagate silencing across the X chromosome. Cell 184, 6174–6192 e6132 (2021). 10.1016/j.cell.2021.10.022

28 Cheng, M. I. et al. Cutting Edge: Hypoxia Sensing by the Histone Demethylase UTX (KDM6A) Limits Colitogenic CD4+ T Cells in Mucosal Inflammation. J Immunol 212, 1069–1074 (2024). 10.4049/jimmunol.2300550

29 Reynolds, S. R. et al. DNA methylation-based immune cell profiling in mouse blood using MouseRS-CMD. Genome Biol 27 (2026). 10.1186/s13059-026-04121-y

30 Yang, T., Ou, J. & Yildirim, E. Xist exerts gene-specific silencing during XCI maintenance and impacts lineage-specific cell differentiation and proliferation during hematopoiesis. Nat Commun 13, 4464 (2022). 10.1038/s41467-022-32273-5

31 Boyman, O., Létourneau, S., Krieg, C. & Sprent, J. Homeostatic proliferation and survival of naïve and memory T cells. Eur J Immunol 39, 2088–2094 (2009). 10.1002/eji.200939444

32 Hickman, S. P. & Turka, L. A. Homeostatic T cell proliferation as a barrier to T cell tolerance. Philos Trans R Soc Lond B Biol Sci 360, 1713–1721 (2005). 10.1098/rstb.2005.1699

33 King, C., Ilic, A., Koelsch, K. & Sarvetnick, N. Homeostatic expansion of T cells during immune insufficiency generates autoimmunity. Cell 117, 265–277 (2004). 10.1016/s0092-8674(04)00335-6

34 Garrett, W. S., Gordon, J. I. & Glimcher, L. H. Homeostasis and inflammation in the intestine. Cell 140, 859–870 (2010). 10.1016/j.cell.2010.01.023

35 Kelsen, J. R. & Sullivan, K. E. Inflammatory Bowel Disease in Primary Immunodeficiencies. Curr Allergy Asthma Rep 17, 57 (2017). 10.1007/s11882-017-0724-z

36 Tegtmeyer, D., Seidl, M., Gerner, P., Baumann, U. & Klemann, C. Inflammatory bowel disease caused by primary immunodeficiencies-Clinical presentations, review of literature, and proposal of a rational diagnostic algorithm. Pediatr Allergy Immunol 28, 412–429 (2017). 10.1111/pai.12734

37 Shi, J. & Petrie, H. T. Activation kinetics and off-target effects of thymus-initiated cre transgenes. PLoS One 7, e46590 (2012). 10.1371/journal.pone.0046590

38 Lovell, C. D. et al. Female mice with a Xist deletion in B cells can develop lupus-associated phenotypes. Cell Rep 45, 117517 (2026). 10.1016/j.celrep.2026.117517

39 Savill, J. & Fadok, V. Corpse clearance defines the meaning of cell death. Nature 407, 784–788 (2000). 10.1038/35037722

40 Sin, J. H. et al. The epigenetic regulator ATF7ip inhibits Il2 expression, regulating Th17 responses. J Exp Med 216, 2024–2037 (2019). 10.1084/jem.20182316

41 Engel, P., Eck, M. J. & Terhorst, C. The SAP and SLAM families in immune responses and X-linked lymphoproliferative disease. Nature Reviews Immunology 3, 813–821 (2003). 10.1038/nri1202

42 Snow, A. L. et al. Restimulation-induced apoptosis of T cells is impaired in patients with X-linked lymphoproliferative disease caused by SAP deficiency. J Clin Invest 119, 2976–2989 (2009). 10.1172/jci39518

43 Moran, A. E. et al. T cell receptor signal strength in Treg and iNKT cell development demonstrated by a novel fluorescent reporter mouse. J Exp Med 208, 1279–1289 (2011). 10.1084/jem.20110308

44 Gaud, G., Lesourne, R. & Love, P. E. Regulatory mechanisms in T cell receptor signalling. Nature Reviews Immunology 18, 485–497 (2018). 10.1038/s41577-018-0020-8

45 Kyle, J. Crohn’s disease in the northeastern and northern Isles of Scotland: an epidemiological review. Gastroenterology 103, 392–399 (1992). 10.1016/0016-5085(92)90826-k

46 Latour, P., Louis, E. & Belaiche, J. Incidence of inflammatory bowel disease in the area of Liege: a 3 years prospective study (1993-1996). Acta Gastroenterol Belg 61, 410–413 (1998).

47 Shivananda, S. et al. Epidemiology of Crohn’s disease in Regio Leiden, The Netherlands. A population study from 1979 to 1983. Gastroenterology 93, 966-974 (1987). 10.1016/0016-5085(87)90558-0

48 McGovern, D. P., Kugathasan, S. & Cho, J. H. Genetics of Inflammatory Bowel Diseases. Gastroenterology 149, 1163–1176 e1162 (2015). 10.1053/j.gastro.2015.08.001

49 Liu, J. Z. et al. Association analyses identify 38 susceptibility loci for inflammatory bowel disease and highlight shared genetic risk across populations. Nat Genet 47, 979–986 (2015). 10.1038/ng.3359

50 Mottet, C., Uhlig, H. H. & Powrie, F. Cutting edge: cure of colitis by CD4+CD25+ regulatory T cells. J Immunol 170, 3939–3943 (2003). 10.4049/jimmunol.170.8.3939

51 Kong, L. et al. The landscape of immune dysregulation in Crohn’s disease revealed through single-cell transcriptomic profiling in the ileum and colon. Immunity 56, 444–458 e445 (2023). 10.1016/j.immuni.2023.01.002

52 Yang, W. & Cong, Y. Exploring Colitis through Dynamic T Cell Adoptive Transfer Models. Inflamm Bowel Dis 29, 1673–1680 (2023). 10.1093/ibd/izad160

53 Wilkinson, N. M., Chen, H. C., Lechner, M. G. & Su, M. A. Sex Differences in Immunity. Annu Rev Immunol 40, 75–94 (2022). 10.1146/annurev-immunol-101320-125133

54 Lee, J. & Chang, H. Y. XIST RNA-protein complex in female-biased autoimmunity: From molecular scaffolds to new clinical biomarkers. J Invest Dermatol 146, 2360–2363 (2026). 10.1016/j.jid.2026.04.018

55 Chen, G. et al. Increased proliferation of CD8+ T cells in SAP-deficient mice is associated with impaired activation-induced cell death. Eur J Immunol 37, 663–674 (2007). 10.1002/eji.200636417

56 Gaspar, H. B., Sharifi, R., Gilmour, K. C. & Thrasher, A. J. X-linked lymphoproliferative disease: clinical, diagnostic and molecular perspective. Br J Haematol 119, 585–595 (2002). 10.1046/j.1365-2141.2002.03851.x

57 Berletch, J. B., Yang, F., Xu, J., Carrel, L. & Disteche, C. M. Genes that escape from X inactivation. Hum Genet 130, 237–245 (2011). 10.1007/s00439-011-1011-z

58 Haribhai, D. et al. A central role for induced regulatory T cells in tolerance induction in experimental colitis. J Immunol 182, 3461–3468 (2009). 10.4049/jimmunol.0802535

59 Steinbach, E. C., Gipson, G. R. & Sheikh, S. Z. Induction of Murine Intestinal Inflammation by Adoptive Transfer of Effector CD4+ CD45RB high T Cells into Immunodeficient Mice. J Vis Exp (2015). 10.3791/52533

60 Erben, U. et al. A guide to histomorphological evaluation of intestinal inflammation in mouse models. Int J Clin Exp Pathol 7, 4557–4576 (2014).

